# Nucleosome remodelling at origins of Global Genome-Nucleotide Excision Repair occurs at the boundaries of higher-order chromatin structure

**DOI:** 10.1101/283747

**Authors:** Patrick van Eijk, Shuvro Prokash Nandi, Shirong Yu, Mark Bennett, Matthew Leadbitter, Yumin Teng, Simon H. Reed

**Author notes:** To whom correspondence should be addressed, Institute of Cancer & Genetics School of Medicine Cardiff University, Heath Park, Cardiff, CF14 4XN, UK.

## Abstract

Repair of UV-induced DNA damage requires chromatin remodeling. How repair is initiated in chromatin remains largely unknown. We recently demonstrated that Global Genome Nucleotide Excision Repair (GG-NER) in chromatin is organized into domains around open reading frames. Here, we identify these domains, and by examining DNA damage-induced changes in the linear structure of nucleosomes, we demonstrate how chromatin remodeling is initiated during repair. In undamaged cells, we show that the GG-NER complex occupies chromatin at nucleosome free regions of specific gene promoters. This establishes the nucleosome structure at these genomic locations, which we refer to as GG-NER complex binding sites (GCBS’s). We demonstrate that these sites are frequently located at genomic boundaries that delineate chromasomally interacting domains (CIDs). These boundaries define domains of higher-order nucleosome-nucleosome interaction. We show that efficient repair of DNA damage in chromatin is initiated following disruption of H2A.Z-containing nucleosomes adjacent to GCBSs by the GG-NER complex.

## Introduction

The basic unit of primary chromatin structure is known as the nucleosome. It comprises a histone octamer, containing two copies each of the canonical four core histones, which are enveloped by the winding of 147bp of DNA around the octamer. The chemical properties of histones can be altered following the post-translational modification of their N-terminal tails. In addition to the canonical histones H2A, H2B, H3 and H4, histone variants such as H2A.Z and H3.3 also exist (Gurard-Levin et al. 2014). Decoration of histone tails with chemical moieties such as ubiquitin, methyl and acetyl groups, or indeed the exchange of histone variants within the octamer structure, alters the physico-chemical properties of the nucleosome, imbuing it with biological information. It is known that the physical arrangement of the nucleosomes in the genome provides an important framework that supports the ordered modification of histone tails and variant turnover. This complex mechanism is known to be important in regulating many chromatin-related biological functions of the cell, including DNA replication, transcription and DNA repair (Lai and Pugh 2017). Defects in such regulatory processes are also implicated in diseases associated with ageing, including cancer (Luijsterburg and van Attikum 2011).

Several decades of research into the fundamental biochemical mechanisms of DNA repair have revealed the basic functions of the multiple pathways that evolved in cells to recognise, remove and correct a bewildering variety of lesions that frequently occur in the genomes of cells (Friedberg et al. 1995). Such damage can be caused by influences of both the internal environment of cells, as well as the broader, external environment in which they exist (Lindahl 1993). One of the major DNA repair pathways is known as nucleotide excision repair (NER) and a great deal is known about its fundamental molecular mechanism (Friedberg 2003; Marteijn et al. 2014). Damaged DNA is excised from the genome as an oligonucleotide of around 30 nucleotides in length by one of two sub-pathways; transcription-coupled NER (TC-NER), and global genome NER (GG-NER). The two pathways differ in the way in which DNA repair is initiated. In TC-NER, this is achieved following the recognition of damage-stalled RNA polymerase II, as it encounters, and is halted by DNA damage. This occurs only in the transcribed strand of actively transcribing genes where DNA damage impedes the progress of RNA polymerase II during transcription. Specific factors exist that mediate this mechanism, which manifests as a more rapid rate of removal of DNA damage from the transcribed strand of active genes compared with other regions of the genome. During GG-NER, which operates in all non-transcribing regions of the genome, including the transcribed strand of silent genes, repair is initiated by a different mechanism, involving a different set of DNA repair factors that function uniquely in GG-NER (Verhage et al. 1994; Reed et al. 1999).

Recent structural and biophysical studies have provided remarkable insight into the mechanism that promotes the recognition of a broad range of different types of damaged DNA bases by GG-NER (Marteijn et al. 2014). These studies helped to define a novel, so-called promiscuous damage recognition mechanism, in which the conformation of the sugar-phosphate backbone of DNA is probed to detect alterations to its structure caused by DNA damage, as opposed to detecting the chemically diverse nature of the lesions themselves (Min and Pavletich 2007; Roche et al. 2008; Sugasawa et al. 2009). Progress is being made with regard to discovering the histone modifications necessary to permit the efficient repair of DNA damage in chromatin, including those modifications that promote NER. A broad range of research has revealed a role for histone ubiquitylation, methylation, acetylation, and histone variant exchange in a variety of DNA repair pathways (Adam et al. 2015; Polo 2015; Polo and Almouzni 2015). However, at this stage, the precise details of how these modifications enhance repair of damage from chromatin remains to be determined.

We previously purified a complex of proteins from yeast cells that is uniquely required for GG-NER in this simple eukaryote (Reed et al. 1999). The complex is comprised of the SWI/SNF superfamily member, Rad16, the Rad7 protein and the yeast general regulatory factor Abf1. We later demonstrated that this complex interacts with the cullin, Cul3 and the elongin, Elc1, to form a UV-inducible E3 ubiquitin ligase that is required for efficient NER (Gillette et al. 2006). We demonstrated that the GG-NER complex regulates UV-induced histone H3 acetylation, by controlling chromatin occupancy of the histone acetyl transferase, Gcn5 at this locus (Teng et al. 2008). This UV-induced hyperacetylation of histones promotes an open chromatin conformation required for efficient repair of DNA damage (Yu et al. 2005; Teng et al. 2008).

We’ve developed genomic tools for the analysis of DNA damage and repair (Teng et al. 2011; Powell et al. 2015) (Yu et al. 2016). We showed that enhanced DNA repair rates occur within open reading frames; a result of the concerted action of both the TC-NER and GG-NER pathways in these regions. Inactivation of the GG-NER complex by deletion of the *RAD7* or *RAD16* genes, resulted in a striking alteration to the genomic distribution of DNA repair rates (Yu et al. 2016), suggesting the presence of repair domains. In the present study, we examined these domains in greater detail to define them, and to determine how chromatin structure is altered by the GG-NER pathway to promote DNA repair at the level the nucleosome, its primary structural unit. To investigate this, we digested chromatin with micrococcal nuclease (MNase), which preferentially digests linker DNA, leaving core nucleosomal DNA largely intact. Coupling the digestion of chromatin with high throughput sequencing is a method referred to as MNase-seq (Jiang and Pugh 2009; Zhang and Pugh 2011). The method enables mapping of nucleosomes across the entire genome, revealing their physical organisation. Two key factors determine the physical arrangement of nucleosomes in the genome. Cis-acting factors include DNA sequence elements themselves, but the effects of these on nucleosome structure can be modified by trans-acting regulatory factors that are activated in response to changes in environmental conditions. These include the exposure of cells to DNA damage induced by UV radiation. Dynamic nucleosomes are typically associated with functional responses, such as regulation of gene expression in the cell to environmental changes or stress (Xi et al. 2011). Changes to the physical organisation of chromatin controls the accessibility of DNA to binding proteins such as transcription factors, replication factors and DNA repair complexes, thereby regulating these activities in the cell. Therefore, using a combination of MNase-seq, ChIP-seq and 3D-DIP-Chip, here we identify the genomic distribution of GG-NER complex chromatin occupancy, and define these positions as GG-NER complex binding sites (GCBS’s). We show that in undamaged cells the GG-NER complex occupies certain gene promoter, nucleosome free regions (NFRs).

Having established the genomic locations of GCBS’s, we observed that they occupy newly-identified boundary sites found throughout the genome. These boundaries delineate chromosomally interacting domains (CIDs) in the genome (Hsieh et al. 2015), which define domains of higher-order nucleosome-nucleosome interaction (Hsieh et al. 2015; Hsieh et al. 2016). Our observations show that GG-NER is organised and initiated from these genomic features. In response to UV damage, we demonstrate that the GG-NER complex remodels H2A.Z-containing nucleosomes immediately adjacent to GCBS’s, to promote efficient DNA repair. This permits redistribution of GG-NER complex components to neighbouring regions of chromatin, where they promote efficient repair of DNA damage. This remodelling process depends on the GG-NER complex, since its absence results in a significant reduction of DNA repair rates in the surrounding regions of the genome. We discuss a paradigm by which the organisation of GG-NER into higher-order chromatin domains, dramatically reduces the genomic search space for DNA damage detection, thus promoting the efficient recognition and repair of DNA lesions throughout the genome.

## Results

### Identification of the genome-wide UV-induced changes to the linear arrangement of nucleosomes

The physical arrangement of nucleosomes can be thought of as a structured array of nucleosome units distributed throughout the genome. Within a population of cells, the precise translational setting of a nucleosome within its unit, in any given cell, may vary centering at a favoured site, which is commonly referred to as its nucleosome position. A single nucleosome position and its change in response to environmental conditions can be characterised by a combination of three parameters that define it. Firstly, the nucleosome position, secondly, its occupancy and finally, its fuzziness, with the latter term meaning the degree of freedom that a nucleosome has, to take up its unitary position within in a population of cells. This degree of freedom is high when a fuzzy nucleosome takes up a wider range of positions in a cell population, and *vice versa* for low fuzziness nucleosomes. In addition to describing the position and fuzziness score of a nucleosome unit, it is also possible to measure its occupancy, which is defined by its peak height as shown in Figure 1A. This refers to the frequency that the nucleosome unit is occupied by nucleosomes within the population of cells.

**Figure 1.**
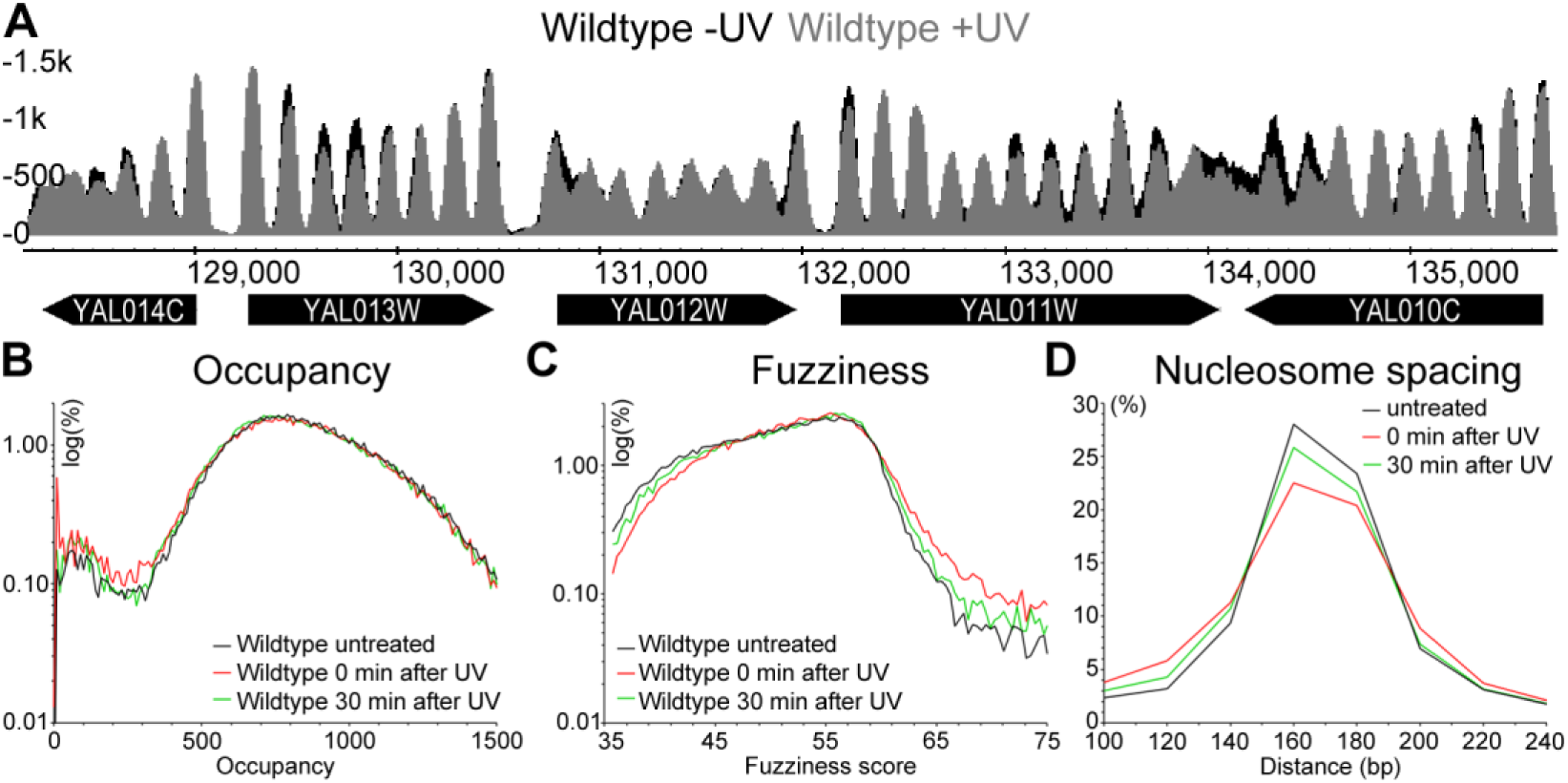
UV-induced changes to the genome-wide nucleosome landscape. A) Represented here are the nucleosomes traces of wild type cells before (black) and after UV irradiation (grey) of an 8 Kbp region on chromosome I (128,000 to 136,000). The genes and their systemic names are indicated by the black arrows underneath the traces. The y-axis on the left indicate the relative read-count that defines the nucleosome peaks in this regions. B) Genome-wide changes to wild type nucleosome occupancy (peak height) in response to UV irradiation are quantified here. The distribution of relative occupancy (in reads) of all >60,000 nucleosomes as a log-scale of percentage is shown here. C) As B but now quantifying the changes to Fuzziness of all nucleosomes in response to UV irradiation. D) As B and C but now quantifying the change in nucleosome spacing of all nucleosomes after UV irradiation.

Trans-acting factors can alter nucleosome structure by changing the position and/or fuzziness of a nucleosome, as well as affecting the nucleosome occupancy at any given position in response to environmental changes. Consequently, we used MNase-seq to map nucleosomes and measure alterations to their structure in wildtype cells exposed to 100 J/m^2^ of UV irradiation. We then compared the changes in nucleosome structure in untreated versus treated cells at different times after UV irradiation, using a bioinformatics pipeline known as DANPOS. This software was specifically developed for measuring genomic changes in nucleosome position, fuzziness and occupancy in cells under different environmental conditions (Chen et al. 2013). In this pipeline, we consistently mapped >60,000 nucleosome positions with high accuracy using this method. In response to UV irradiation, changes to nucleosome occupancy at various positions across the genome are readily observed when the mapped nucleosome traces are plotted in a linear fashion. A representative 8 Kbp section of yeast chromosome I is shown in Figure 1A (note that occupancies with or without UV exposure of cells are indicated in black and grey, respectively). The aggregate changes in nucleosome occupancy, fuzziness and position for all >60,000 nucleosomes are summarised in Figure 1B to D. Here, major changes to nucleosome occupancy throughout the genome are not observed (Figure 1B). This indicates that wholesale loss of nucleosome structure does not occur in response to UV damage. A small accumulation of nucleosomes at sites of low-occupancy nucleosomes (<350 normalised reads) can be detected immediately after UV irradiation (Figure 1B, red line). Genome-wide nucleosomes fuzziness of nucleosomes on the other hand, is altered to a greater extent in response to UV. Thus, we observed an increase in the number of fuzzy nucleosomes detected, with a reciprocal decrease in low-fuzziness nucleosomes that were initially well-positioned, at both 0 and 30 minutes after UV irradiation (Figure 1C). Finally, we express the genomic position of nucleosomes as the internucleosomal distance or nucleosome spacing in base pairs. As expected, the average spacing for all nucleosomes, as defined by the length of linker and nucleosomal DNA, is enriched for distances of between 160 to 180 base pairs, as shown in Figure 1D. As a result of UV irradiation, a small loss of nucleosomes with this spacing is observed, with a reciprocal gain in more closely spaced nucleosomes (i.e. those with <140bp spacing). Our observations thereby identify the UV-induced changes to the linear nucleosome structure throughout the entire genome, revealing that chromatin is remodelled at only a sub-set of positions, via discrete local changes dispersed throughout the genome. Since UV-induced lesions are essentially distributed uniformly throughout the genome, our observations suggest that repair of damage may be initiated through nucleosome remodelling at a limited number of specific nucleosomal sites in the genome.

### The canonical nucleosome structure at transcription start sites is maintained following UV exposure

We considered whether the UV-induced changes in nucleosome structure described above can be observed in the typical phasing of nucleosomes found at transcription start sites (TSS’s). Therefore, we mapped nucleosome positions in relation to annotated TSS’s (Xu et al. 2009). We find that, when all 5,171 TSSs are examined, the nucleosome structure around these genomic features remains essentially unaltered after UV irradiation (A). Nucleosome structure around TSSs is consistently made up of a pronounced nucleosome free (NFR) or depleted (NDR) region just upstream of the TSS, flanked by a well-defined +1 nucleosome overlying the TSS. This is followed by an array of positioned nucleosomes extending into more distal regions of the gene’s open reading frame (ORF). Upstream of the NFR, another array of more fuzzy and lower occupancy nucleosomes can be discerned (Figure 2A), as described previously (Jiang and Pugh 2009). This result demonstrates that no gross UV-induced changes to nucleosome structure occur in the context of this well-established genomic feature of nucleosome organisation. Disruption of nucleosome structure at this feature has been previously described for mutants defective in certain essential, SWI/SNF ATP-dependent chromatin remodellers, including *CHD1, ISW1* or *INO80* (van Bakel et al. 2013), which regulate gene expression by controlling nucleosome structure. Indeed, in yeast, nucleosome sliding, which shifts the translational setting of the nucleosome, altering its position, is a well-known mechanism to control gene expression (van Bakel et al. 2013). Part of the cellular response to DNA damage controls the gene expression of various DNA-damage responsive genes via this mechanism. Therefore, we investigated nucleosome sliding at a single UV-responsive gene by plotting nucleosomes at the DNA damage inducible locus, *RAD51* (Shinohara et al. 1992), as shown in Figure 2B. As expected, these data demonstrate that nucleosome sliding can be detected at this locus after DNA damage induction. The absence of linear nucleosome sliding when *all* TSSs are examined in aggregate, however, indicates that this mechanism of nucleosome remodelling does not occur widely throughout the genome in response to UV damage. Collectively, these results indicate that the UV-induced nucleosome remodelling observed in Figure 1 occurs in a different genomic context.

**Figure 2.**
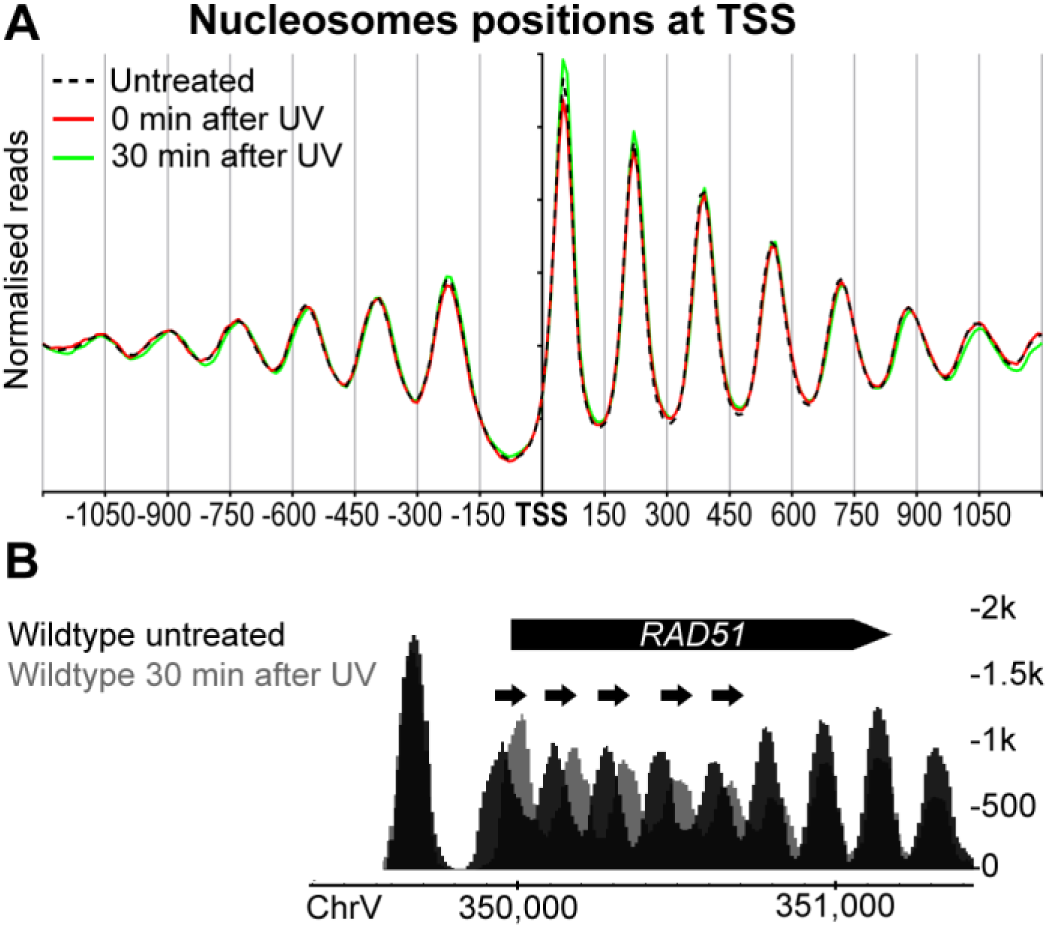
Nucleosomes occupancy around *all* TSS in yeast does not change in response to UV irradiation in wild type cells. A) Composite plot of nucleosomes positions relative to all TSS (n = 5,171). Genome-wide MNase-seq data was used to aggregate nucleosome positioning in relation to TSS positions. B) Genome browser snapshot of the DNA damage inducible gene *RAD51* (chromosome V) displaying the nucleosome positions prior to DNA damage (black) and 30 minutes after UV irradiation (grey). The black arrows indicate the shift of nucleosomes. The black bar represents the ORF. The y-axis on the right indicate the relative occupancy of the nucleosomes in reads.

### Identifying the chromatin context for the linear remodelling of nucleosomes during GG-NER

In order to identify sites of chromatin remodelling at the level of individual nucleosomes during GG-NER, we measured nucleosome positions in relation to GG-NER complex binding in chromatin. We recently reported that GG-NER is organised into domains defined by the chromatin occupancy of the Abf1 component of the GG-NER complex. Abf1 is a general regulatory factor that functions in concert with chromatin remodellers to establish the basic pattern of linear nucleosome structure at the 5’ ends of genes, resulting in the formation of chromatin barrier structures within the genome (Krietenstein et al. 2016). Therefore, we performed an Abf1 ChIP-seq experiment, mapping the precise genome-wide occupancy of Abf1 to chromatin at nucleotide resolution as a precursor to identifying locations of GG-NER complex binding. This enabled us to accurately map the nucleosome structure at positions of GG-NER complex binding in a similar fashion to that shown in Figure 2A. In agreement with our previously published Abf1 ChIP-chip data (Yu et al. 2016), we found ~4,000 Abf1 binding sites detected by MACS2 (data not shown). It is well established that Abf1 exists in the cell in large excess over the other GG-NER components (Rad7 and Rad16), and has a wide range of different functions outside of GG-NER (Yarragudi et al. 2007; Schlecht et al. 2008; Ganapathi et al. 2011; Zhang et al. 2012). We sub-classified these Abf1 binding sites by identifying those that overlap, either with Abf1 consensus binding sequences distributed throughout the yeast genome (n = 1,752), or a list of curated genome-wide NFRs (n = 6,589), at which Abf1 can frequently be found (Figure 3A) (Yarragudi et al. 2007; Hartley and Madhani 2009; Ganapathi et al. 2011; Ozonov and van Nimwegen 2013). The Venn-diagram in Figure 3A enables us to identify various subcategories of Abf1 binding sites. We then used our previously published Rad16 ChIP-chip genome-wide occupancy data (Yu et al. 2016) to refine this list by identifying the genomic positions enriched for GG-NER complex binding. This yielded 3,600 Abf1 binding sites that are enriched for Rad16. The majority of these genomic positions (~70%) are located upstream of genes, with only 13% of these positions mapping within ORFs, and another 10% found downstream of genes. We used this information to further refine the set of GG-NER complex binding positions among them. This set of upstream Abf1 binding positions (n = 2664 i.e. ~70%) we now refer to as GG-NER Complex Binding Sites (GCBS’s). Using this list, we generated composite plots of the nucleosome positions at *all* GCBS’s. As expected, the high percentage of overlap between NFRs and GG-NER complex binding sites reveals a nucleosome-depleted region at these genomic positions (Figure 3B, black line). These sites are flanked by an array of positioned nucleosomes. Next, we investigated the effect of UV irradiation on nucleosome structure at these positions. In wild type cells, loss of nucleosome occupancy at GCBS-adjacent nucleosomes can be discerned immediately after UV irradiation (Figure 3B, red line). After 30 minutes of repair time, nucleosome occupancy returns to normal levels, with some increased fuzziness (Figure 3B, green line). This identifies the genomic location of remodeled nucleosomes in response to UV radiation.

**Figure 3.**
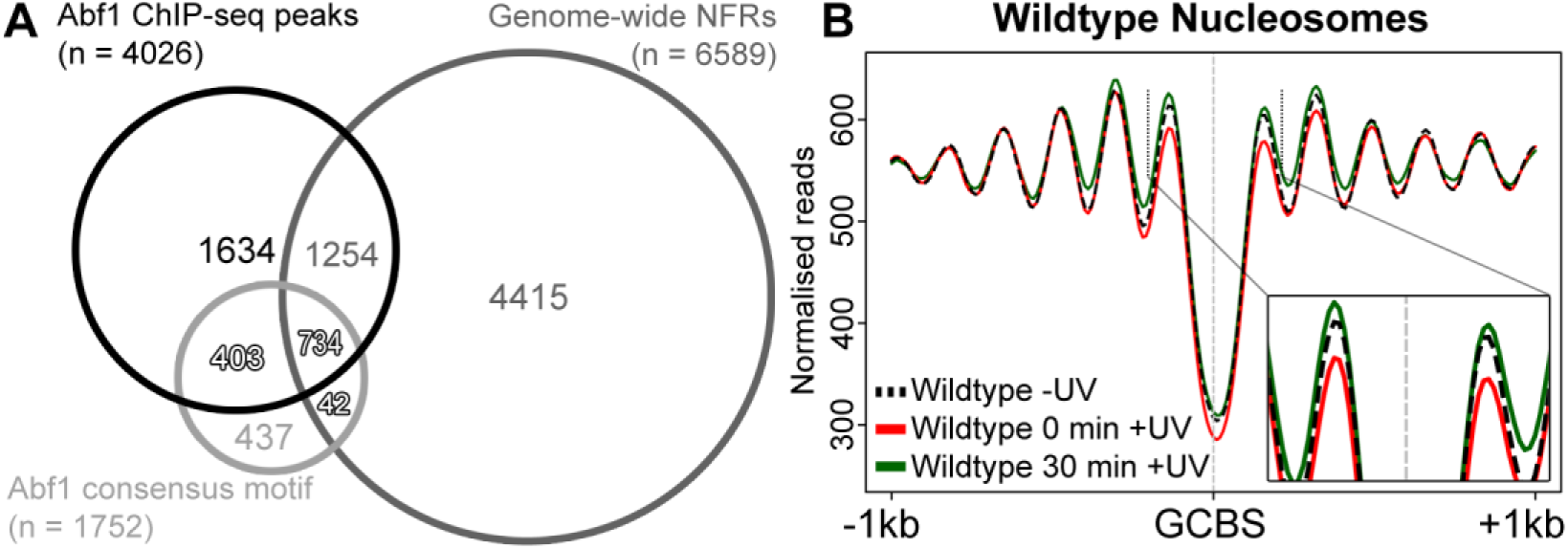
GG-NER complex binding sites are positions of nucleosome organisation and UV-induced remodelling. A) The Venn diagram presented here subcatergorises the ChIP-seq Abf1 binding sites into classes based on their localisation to NFRs (n = 6589) and/or Abf1 consensus motifs (n = 1752) used to identify GCBS’s. The numbers indicate the amount of overlapping events of that category allowing no gap between the features. B) MNase-seq data of wild type cells was used to plot cumulative nucleosome positions around GCBS’s (n = 2664) in the absence of UV irradiation and at different intervals after UV irradiation, displaying regularly spaced nucleosome arrays at these genomic locations. The x-axis denotes the 2 Kbp regions surrounding the GCBS’s while the y-axis indicates nucleosome occupancy as measured by normalised reads.

### GG-NER complex binding sites are located at the boundary regions of Chromosomally Interacting Domains

We detected UV-induced nucleosome remodelling in only a small subset of nucleosomes (Figure 3B). However, standard MNase-seq can only reveal changes in the linear arrangement of nucleosomes. Therefore, we investigated the genomic locations of GCBS’s in relation to domains of higher-order chromatin structure. Recent advances in methods such as 3C and the related HiC methods, have led to the introduction of a chromatin capture method called Micro-C (Hsieh et al. 2015; Hsieh et al. 2016). This method measures higher-order nucleosome-nucleosome interactions in chromatin. Micro-C follows the same principles as other 3C methods, but uses MNase instead of restriction enzymes to digest cross-linked chromatin. This allows the detection of distal nucleosome-nucleosome interactions that have recently led to the discovery of chromosomally interacting domains (CIDs) at nucleosome resolution for the first time in yeast (Hsieh et al. 2015). The boundary sites that demarcate CIDs are often found upstream of highly expressed genes, and are enriched for nucleosome pairs that flank NFRs. Therefore, we set out to determine the relationship between the genomic locations of the GCBS’s that we identified above, and the boundary sites of these newly discovered CIDs. To this end, we retrieved the genomic positions of the CID boundaries from published data (Hsieh et al. 2015), and calculated the overlap between these positions and the GCBS’s. We find that GCBS’s map predominantly to CID boundary positions, with around 50% mapping precisely to these sites, as shown in Figure 4A. To examine the significance of this observation, we took a similar number of random genomic intervals and calculated the overlap of boundaries and GCBS’s with these sites. This revealed only two boundaries overlapping at these positions compared with the 1200 GCBS’s found at CID boundaries (Figure 4A). This confirms that GCBS’s colocalise at the boundary positions of a specific set of CIDs, and therefore occupy regions of the genome that demarcate higher-order nucleosome-nucleosome interactions. Figure 4B shows a representation of the newly discovered CID chromatin landscape in relation to the linear setting of nucleosomes (D), and also shows the position of two GCBS’s at CID boundaries, as illustrated by the binding of Abf1 (C). We conclude that GG-NER is organized and initiated from the boundaries of specific CIDs, to which the complex is bound in the absence of DNA damage.

**Figure 4.**
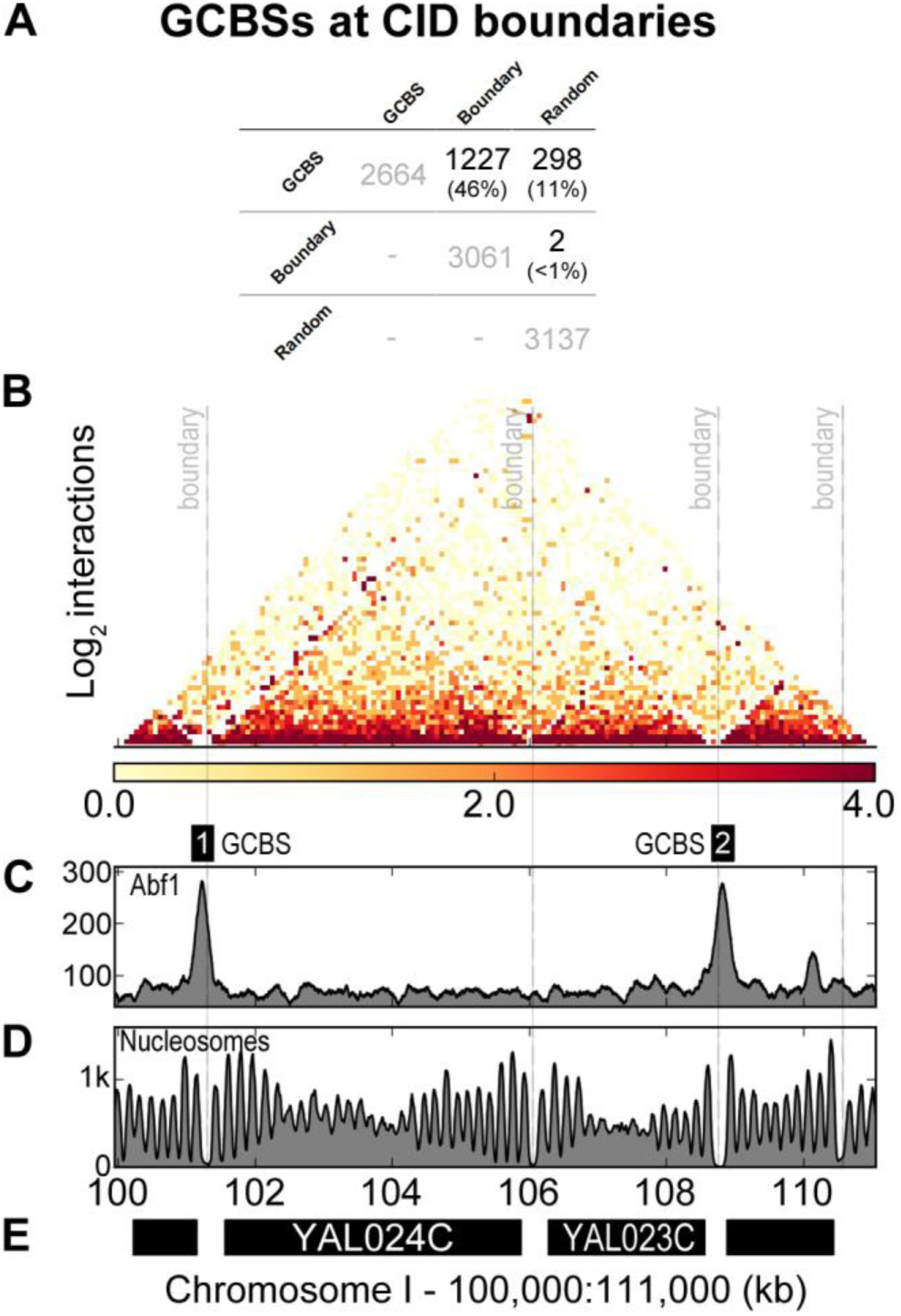
GCBS’s are located at the boundaries of Chromosomally Interacting Domain as characterised by Abf1 binding and NFRs. A) Overlap calculations identified the number and identity of GCBS’s (n = 2664) at boundaries (n = 3061) and at random sites (n = 3137). The percentage of GCBS’s in each subcategory is indicated between brackets. B) Micro-C data (Hsieh et al. 2016) was used to plot nucleosome-nucleosome interactions in a 11Kbp window on chromosome I. The grey dashed lines indicate 4 boundary positions documented in literature (Hsieh et al. 2015). The intensity of the heatmap is a measure for the normalized interactions indicated underneath the panel. C) Abf1 ChIP-seq data is plotted here to highlight two GCBS’s in this region of the genome labelled as GCBS 1 and 2. D) The nucleosome landscape is presented here by plotting MNase-seq data at this genomic interval. E) Indicated in black bars are the genes located in this region of the genome. The x-axis labels highlight the genomic coordinates in Kbp. The y-axis on each panel indicates peak height as normalised reads.

### The GG-NER complex regulates nucleosome structure at GCBS-adjacent nucleosomes

We examined the role of the GG-NER complex on nucleosome structure in the vicinity of GCBS’s, To do this we mapped nucleosomes in *RAD16* deleted cells and compared them to the wild type pattern (A, grey line). We observed reduced nucleosome occupancy at the positions immediately adjacent to the GCBS’s in these cells. This indicates that the GG-NER complex is necessary for establishing the normal nucleosome structure adjacent to these locations in undamaged wildtype cells (Compare black line with grey line, Figure 5A). This demonstrates the importance of the complex in establishing and maintaining nucleosome structure at these sites. Figure 5B demonstrates that in Rad16 deleted cells, no UV-induced loss of nucleosome occupancy is observed at these positions (Figure 5B). This indicates the dependence of the nucleosome remodelling at these positions on the GG-NER complex. In this mutant, nucleosomes actually accumulate 30 minutes after UV DNA damage induction (Figure 5B, green line) compared to the initial levels of nucleosome occupancy at GCBS-adjacent nucleosome positions (Figure 5A). Furthermore, we found no evidence for nucleosome sliding around GCBS’s, which is in agreement with our previous observations that showed that whilst the GG-NER chromatin remodelling complex has DNA translocase activity, it is unable to slide nucleosomes *in vitro* (Yu et al. 2009). The UV-induced loss of nucleosome occupancy observed is consistent with histone exchange events described by others in the context of gene structure (van Bakel et al. 2013). We suggest that the nucleosome remodelling process at the GCBS-adjacent nucleosomes is a mechanism that initiates GG-NER (Weber et al. 2014).

**Figure 5.**
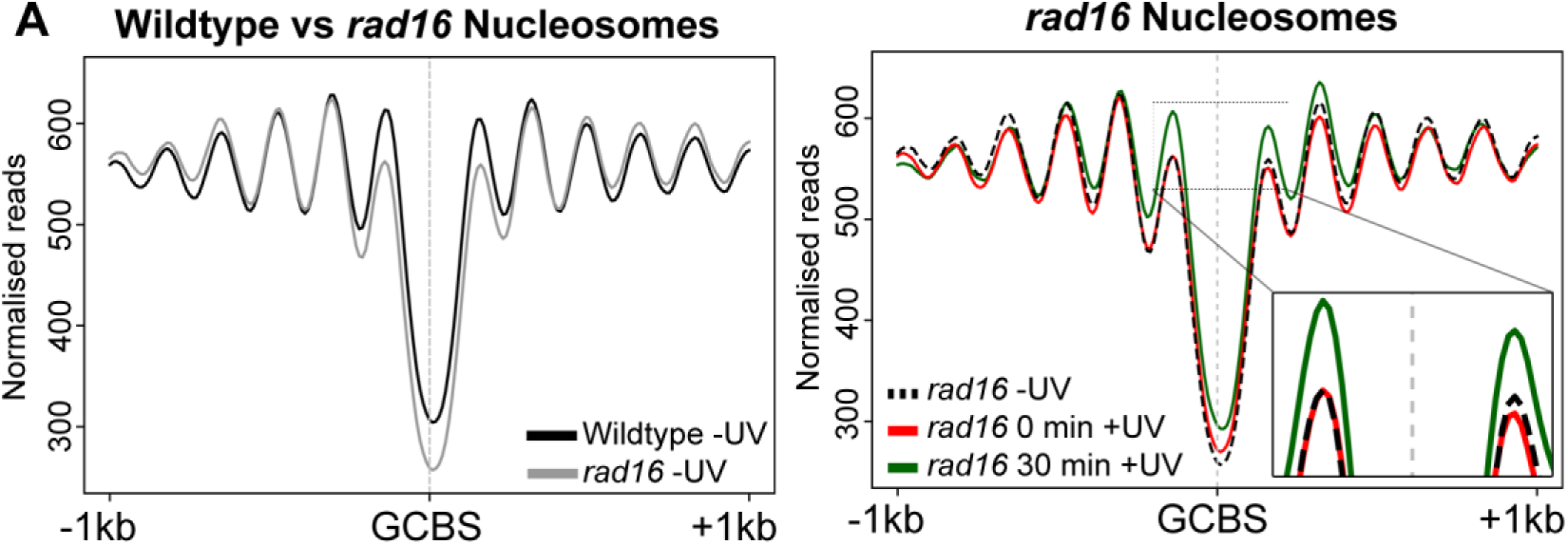
GG-NER complex binding sites are positions of Rad16-dependent nucleosome organisation and UV-induced remodelling. A) MNase-seq data of wild type and *rad16* mutant cells was used to plot cumulative nucleosome positions around GCBS’s (n = 2664) in the absence of UV irradiation, displaying regularly spaced nucleosome arrays at these genomic locations. The x-axis denotes the 2 Kbp regions surrounding the GCBS’s while the y-axis indicates nucleosome occupancy as measured by normalised reads. B) As A, showing UV-induced changes to nucleosomes positions around GCBS’s in *rad16* mutant cells.

### GCBS’s are flanked by histone H2A.Z-containing barrier nucleosomes that are exchanged in response to UV damage

Dynamic nucleosomes are often associated with the functional response of the cell to environmental change or stress (Lai and Pugh 2017). This physical organisation of the chromatin controls the accessibility of binding proteins to the DNA in chromatin, such as transcription factors, thus regulating their activity in the cell. Such nucleosomes contain the histone variant H2A.Z, and have been described previously as ‘barrier nucleosomes’ that represent highly dynamic nucleosome sites in the genome. As such, they represent nodes that are inhibitory structures, which must be altered to permit gene expression at such locations. A role for histone variants in DNA repair has been noted in both NER and other repair mechanisms (Adam et al. 2015). Indeed, in yeast, we have previously reported that histone H2A.Z is involved in NER (Yu et al. 2013). Therefore, we investigated the occupancy of histone H2A.Z at nucleosomes located immediately adjacent to GCBS’s. To do this, we undertook ChIP-seq experiments using HA-tagged H2A.Z to map the positions of genome-wide H2A.Z containing nucleosomes. We then measured the change in their occupancy in response to UV irradiation. When the sequencing data obtained was aligned to the reference genome, and H2A.Z peaks were identified using the DANPOS software package (Chen et al. 2013), we detected about 16,000 H2A.Z containing nucleosomes. Initial analysis of the genome-wide distribution of histone H2A.Z confirmed the presence of this histone variant predominantly at nucleosomes flanking NFRs upstream of genes. These display an asymmetric pattern of binding as described previously in the literature (Guillemette et al. 2005; Raisner et al. 2005; Albert et al. 2007; Weber et al. 2014). Next, we examined the H2A.Z occupancy in GCBS-adjacent nucleosomes and observed H2A.Z at these sites (Figure 6A). Consistent with the loss of nucleosomes observed earlier (Figure 3B), we found that H2A.Z occupancy is also reduced at these positions in response to UV irradiation (Figure 6A, red line). In addition, after 30 minutes of repair time, H2A.Z occupancy returns to pre-damage levels (Figure 6A, green line), which is also consistent with the wild type recovery of nucleosome occupancy shown in Figure 3B.

**Figure 6.**
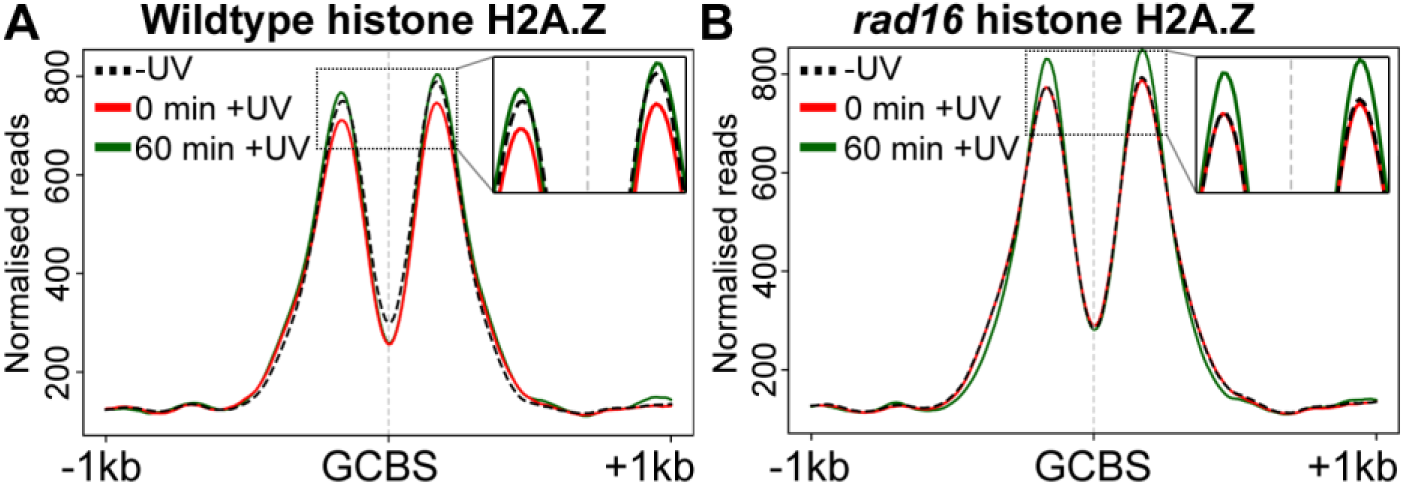
Histone H2A.Z occupancy around GCBS’s changes in response to UV irradiation. A) H2A.Z occupancy adjacent to GCBS’s (n = 2664) is plotted here from ChIP-seq data mapping H2A.Z nucleosomes before and 0 or 60 minutes after UV irradiation. The x-axis represents the GCBS’s and 1 Kbp either side of these positions. H2A.Z occupancy is quantified as normalised ChIP-seq reads on the y-axis. B) As A, plotting H2A.Z nucleosomes in the absence of the *RAD16* GG-NER gene in response to UV irradiation.

We then investigated the GG-NER complex-dependence of H2A.Z eviction at these sites. The histone H2A.Z ChIP-seq experiment was repeated in cells deleted for *RAD16* and the data was processed as described above. In the absence of DNA damage, histone H2A.Z occupies nucleosomes that neighbour the GCBS in *rad16* mutant cells, in a similar fashion to that observed in wildtype cells (compare Figure 6A black dashed line with 6B black dashed line). However, in response to UV, no loss of histone H2A.Z can be detected in the GG-NER defective *RAD16* deleted cells (Figure 6B, red line). Consistent with our findings for nucleosome occupancy (Figure 4B), sixty minutes after UV irradiation, H2A.Z occupancy also increases significantly beyond the level observed prior to DNA damage induction (Figure 6B, green line) to levels higher than that observed in wildtype cells (Figure 6A). Absence of H2A.Z eviction at these sites in a GG-NER defective mutant, confirms a role for the GG-NER complex in this process. These data confirm that histone eviction or exchange of H2A.Z at nucleosomes adjacent to GCBS’s, is driven by the GG-NER complex to alter chromatin structure during the initial stages of GG-NER.

### GG-NER complex remodelling of GCBS-adjacent nucleosomes in relation to gene structure at TSSs

Examining the data in the way described in the previous sections reveals a near symmetrical distribution of nucleosomes and H2A.Z at GCBS’s. This is due to the random orientation of neighbouring ORFs when the data is plotted in this fashion. As shown in Figure 2A, orientating genes according to the direction of transcription, displays a characteristic asymmetry to the nucleosome occupancy in relation to TSSs (Jiang and Pugh 2009). In order to examine GG-NER-dependent nucleosome remodelling in relation to gene orientation, we organised the GCBS’s according to the strand information of the nearest gene they are annotated to, and aligned the peaks to the nearest gene. This orientates the data in such a way that the ORFs are positioned downstream (i.e. to the right) of the GCBS’s, while the GCBS positions that are defined by the position of Abf1 binding, are right-aligned and extend upstream of the ORFs (i.e. to the left). We used this method to display the nucleosome and H2A.Z occupancy data in relation to gene orientation. Plotting nucleosomes and H2A.Z in this way now reveals a nucleosome pattern similar to that shown in Figure 2A (Figure 7A and B). Having identified the precise location of GG-NER complex binding sites in relation to TSSs, we finally examined the UV-induced remodelling of nucleosomes at this subset of genomic locations. We again observed the loss of nucleosome occupancy after UV irradiation at both the −1 and +1 nucleosomes as defined by the NFR (Figure 7A). Therefore, nucleosome eviction or exchange occurs predominantly at these nucleosomes. Moreover, this representation confirms that nucleosome sliding does not occur during GG-NER. We also plotted the H2A.Z data in this context, to examine changes in its occupancy in response to UV irradiation. Figure 6B shows the asymmetric distribution of histone H2A.Z binding around the promoter proximal NFRs. H2A.Z is predominantly found at the +1 nucleosome, which in this class of GCBS-adjacent nucleosomes is located directly at the position of the TSS. Typically, the +1 nucleosome is positioned further into the ORF when nucleosomes are mapped to *all* TSSs (Figure 2A). To determine whether this novel subset of TSS-positioned nucleosomes is unique to this class of GG-NER complex binding sites, we used K-means clustering of the individual nucleosome traces to identify a subclass of genomic positions that uniquely contain a +1 nucleosome at the TSS, or whether this is a common feature amongst these genes. Interestingly, we find 13 clusters that all display a different distance between the TSS and the +1 nucleosome (Supplementary Figure S1A). To represent the nucleosome structure in a different orientation, we used the coordinates of the NFR at these GCBS’s and right-aligned them. Plotting the MNase-seq data in this orientation uniformly aligns the nucleosome arrays revealing the presence of only four clusters (Supplementary Figure S1B). Therefore, we conclude that the +1 nucleosome positions at the TSS as shown in is explained by the averaging of different nucleosome traces that exhibit a highly variable ratio in the distance between the TSS and the +1 nucleosome position, demonstrating that this is not a typical feature of these GCBS-associated regions (Supplementary Figure S1).

**Figure 7.**
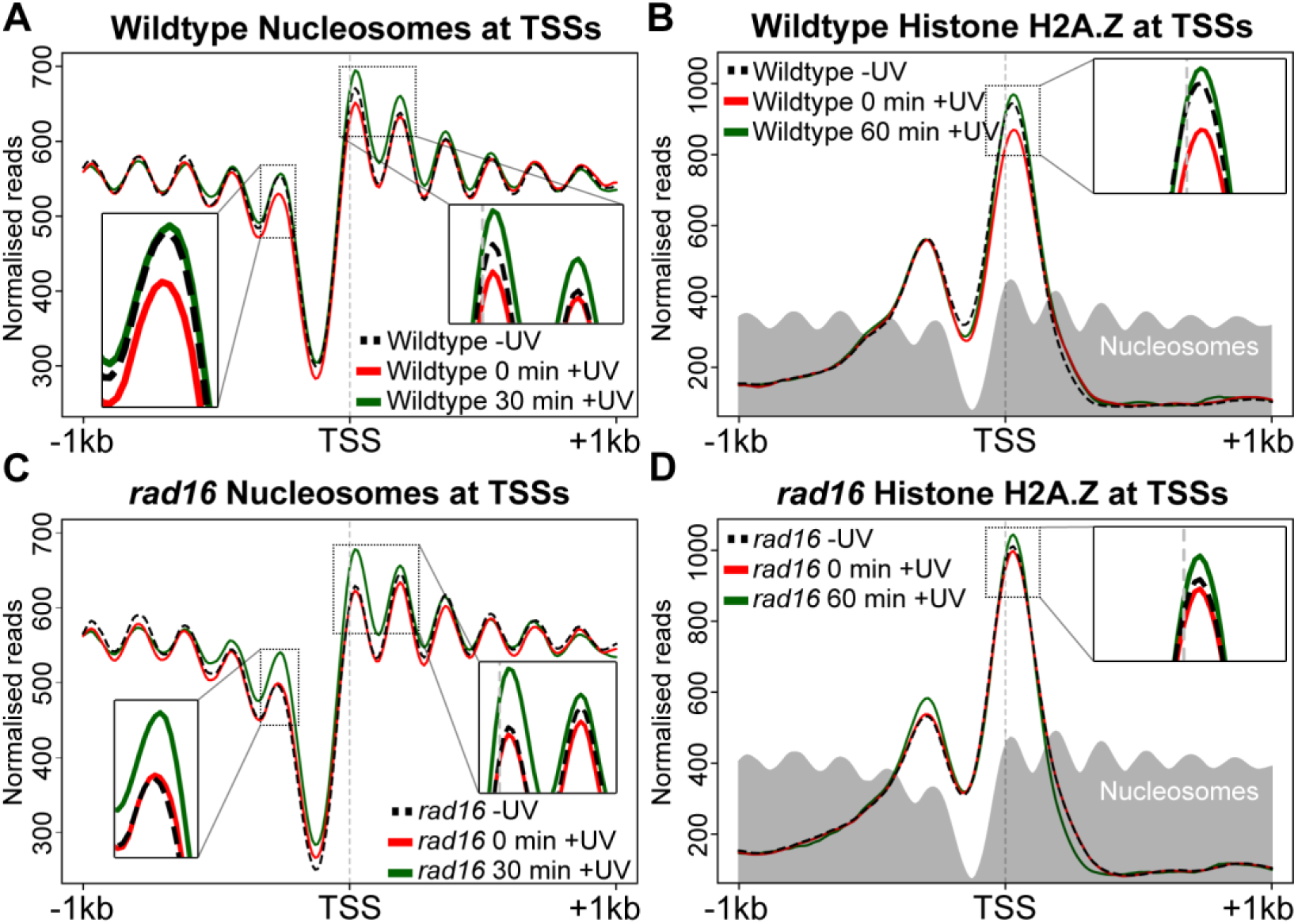
Nucleosome and H2A.Z occupancy around TSS’s of genes that contain a GCBS in their promoter region. A) Nucleosome occupancy in wild type cells is displayed here. MNase-seq data of untreated and UV-treated cells is shown as cumulative graphs around TSS’s (n = 2664). The inserts highlight the nucleosome remodelling at the −1 position (on the left) and the remodelling at positions +1 and +2 (on the right). B) The UV-induced change to H2A.Z occupancy in wild type cells around GCBS-associated TSS’s is shown here using H2A.Z ChIP-seq data prior to UV irradiation and 0 or 60 minutes after UV damage. The light gray trace represents the nucleosome positioning in the absence of DNA damage. The insert focuses on the UV-induced changes to H2A.Z occupancy at the +1 position. C) As A, now representing the nucleosome occupancy at GCBS-bound promoter regions in *RAD16* deleted cells. D) As B, here showing the UV-induced H2A.Z occupancy changes at nucleosomes immediately adjacent to the TSS of GCBS-associated genes in *rad16* mutant cells.

We observed that H2A.Z occupancy at the −1 nucleosome is much lower than at +1 position, and does not change in response to UV irradiation (Figure 7B). However, UV-induced loss of occupancy of H2A.Z at the +1 nucleosomes can be readily detected. H2A.Z occupancy at 60 minutes increases to levels higher than those in untreated cells. These differences may reflect variations in the type and timing of the histone eviction/exchange events occurring in the +1 and −1 nucleosomes during repair. Figure 7C and D demonstrate that the GG-NER complex is required for the eviction or exchange of H2A.Z containing nucleosomes adjacent to the GCBS’s, since loss of nucleosome occupancy is not observed in *RAD16* deleted cells. As previously noted, accumulation of nucleosomes is observed at these sites in repair defective cells. To confirm that this process is specific to GCBS sites, we analysed nucleosomes at all TSSs in the *RAD16* deleted strain and observe no UV-induced changes at those sites (Supplementary Figure S2). Collectively, our data demonstrates that nucleosome remodelling occurs at promoter proximal NFRs at sites of GG-NER complex binding. In response to UV irradiation, histone eviction or exchange occurs at these GCBS-adjacent nucleosomes.

### UV-induced changes in the chromatin occupancy of GG-NER complex components at GCBS’s

We recently demonstrated that the GG-NER complex binds to promoter-proximal regions (Yu et al. 2016), but how its components bind to chromatin in relation to the nucleosome structure that exists at these genomic features is unknown. Therefore, we used the list of annotated TSSs described above and plotted GG-NER complex chromatin occupancy at those features to describe in more detail the GG-NER complex binding at these sites. We used the Abf1 ChIP-seq and Rad16 ChIP-chip data to perform this analysis. As described previously, Abf1 occupancy is highest around the promoter region upstream of the TSS and is located precisely at the NFR (Figure 8A). We also observe a left-sided shoulder to the distribution of Abf1 occupancy upstream of the NFR, which demonstrates that the distribution of NFR sizes to which ABF1 can bind, extends upstream of the TSS. In the absence of UV irradiation, the Rad16 component of the GG-NER complex is enriched at these sites as previously reported.

**Figure 8.**
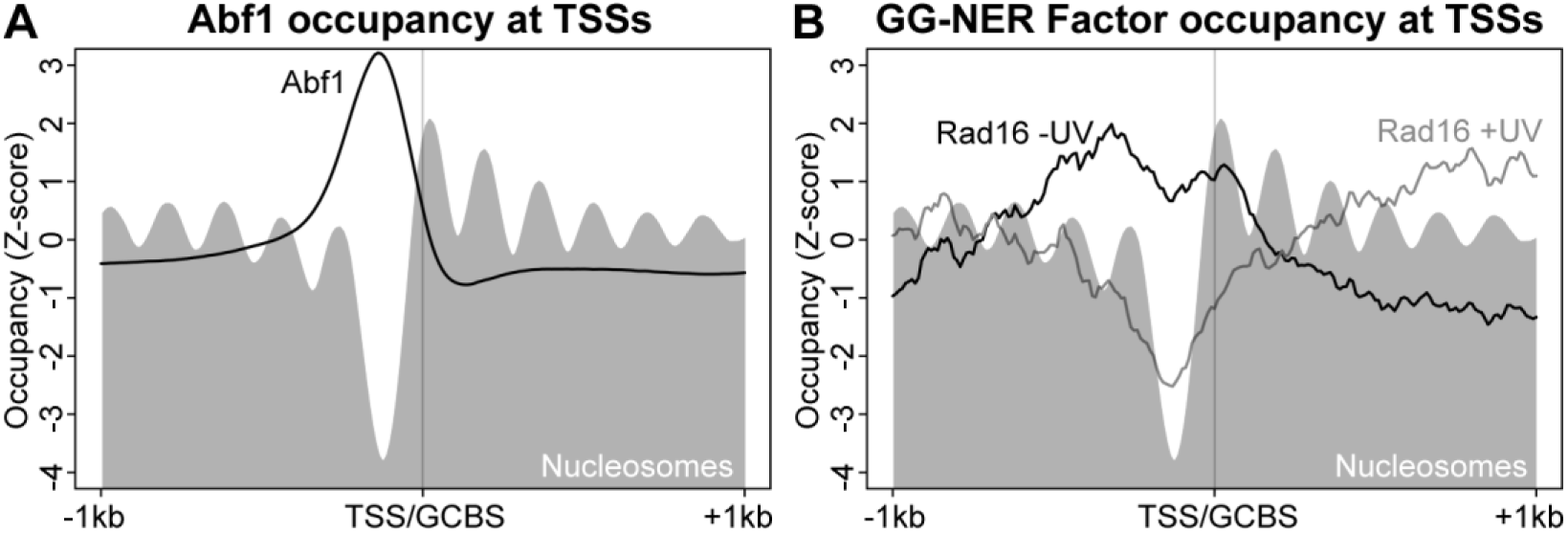
GG-NER complex binding to the chromatin landscape around GCBS-associated promoter regions. A) Genomic positions of GCBS-associated genes were used to plot the genome-wide Abf1 ChIP-seq data at TSS’s to map the location of complex binding in relation to gene structure and nucleosome positions at these genes. The grey nucleosome trace represents nucleosome occupancy in the absence of UV irradiation at these loci. The y-axis represents the Z-score converted reads to enable comparison and plotting of data with different read-depths into the same plot. B) GG-NER complex binding as assessed by Rad16 ChIP-chip data (Yu et al. 2016) plotted at GCBS-associated genes. The grey nucleosome trace represents nucleosome occupancy in the absence of UV irradiation at these loci.

However, when the data is examined in this context, the precise location of Rad16 occupancy can now be discerned. Peaks of Rad16 occupancy are observed predominantly at the +1 and −1 nucleosomes, and Rad16 enrichment also extends to the −2 nucleosome. This is consistent with the upstream shoulder of Abf1 enrichment described above (Figure 8A). Collectively, these data show that the Abf1 component of the GG-NER complex binds to chromatin at specific NFR’s, which are flanked by nucleosomes where peaks of Rad16 binding are observed at the +1 and −1 positions. The occupancy of individual GG-NER complex components at positions upstream of the −1 nucleosome might indicate that Rad16 binds to the −1 nucleosome when the Abf1 component of the complex occupies broader NFRs that are located upstream, and away from the TSS. Indeed, this accounts for the broader range of translational settings observed for Abf1 as shown in Supplementary Figure S2 (right hand panel). In response to UV irradiation, the occupancy of the Rad16 component of the GG-NER complex is rapidly altered from its original position at +1 and −1 promoter nucleosomes, and is redistributed to sites more distally located at nucleosomes extending into the ORF of genes, as well as the upstream regions of promoters. No change in Abf1 occupancy is observed as previously described (Yu et al. 2016).

### GG-NER complex remodelling of nucleosomes at GCBSs establishes domains of efficient GG-NER in the genome

GG-NER is organised from Abf1 binding sites found at intergenic regions around ORFs (Yu et al. 2016). In the current study we have shown that these positions can be refined to reveal how the GG-NER complex promotes nucleosome remodelling. However, until now it has not been possible to map relative rates of DNA repair in relation to nucleosome positions at GCBS’s. Therefore, we plotted 3D-DIP-Chip repair rate data (Yu et al. 2016) at the GCBS’s, where UV-induced nucleosome remodelling occurs, to establish whether these sites are also domains from which repair is organised. In wild type cells, repair rates are evenly distributed across the 2Kbp window surrounding these sites (Figure 9A), with variation in the rates observed in relation to nucleosome positions as reported by others (Mao et al. 2016). However, in the absence of GG-NER, CPD repair rates in the vicinity of GCBS’s are severely reduced, as shown in *RAD16* deleted cells (Figure 9A, grey line). This confirms and extends our previous observations; that the GCBS’s where UV-induced nucleosome remodelling occurs during GG-NER are also the sites from which GG-NER is organised and initiated in the genome.

**Figure 9.**
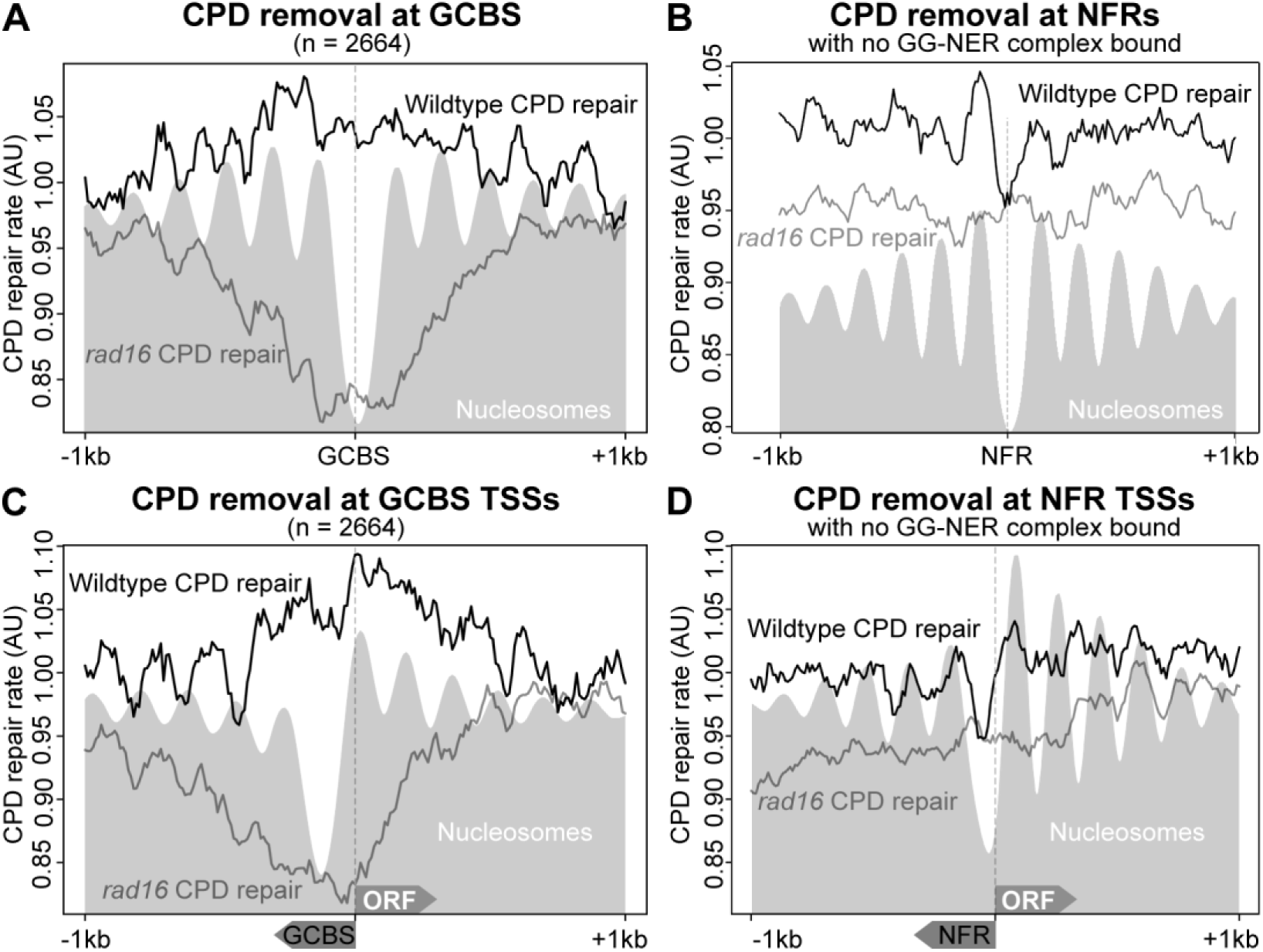
Nucleosome remodelling and repair organisation require chromatin binding of the GG-NER complex and are not common features of NFRs. A) 3D-DIP-Chip data from wild type and GG-NER deficient cells was converted and plotted at GCBS’s to show the organisation or repair as described previously. The nucleosome landscape at these genomic positions is presented as the grey trace, showing the repair rates now in the context of nucleosome positions. The GCBS positions including 1 Kbp on either side are presented here. The CPD repair rates are expressed as arbitrary units. B) As A, now using a set of genome-wide NFRs (n = 4415) to which *no* GG-NER complex is bound, to plot the nucleosome and 3D-DIP-Chip data of wild type and *RAD16* deleted cells. Regions up to 1Kbp up- and downstream of these NFR are used to plot the data. Repair rates are expressed as arbitrary units. C) As A, now using the TSS of the nearest gene to which the GCBS locates. The x-axis indicates the relevant orientation of the GCBS upstream of the TSS and the ORF directed towards the right (downstream). D) As B, here reorienting the data towards the nearest gene and aligning the data at the TSS. The x-axis runs 1 Kbp up- and downstream from these positions. The NFRs are positioned upstream of the TSS, whereas the ORFs are positioned downstream.

The use of genome-wide NFR, nucleosome, and Abf1 ChIP-seq data allowed us to annotate the Abf1 binding sites and identify GCBS’s with nucleotide resolution. This combination of features also enables us to select a set of genomic NFRs that are not associated with GG-NER complex binding. This set of unrelated NFRs can therefore serve as *in silico* controls, since no GG-NER-dependent nucleosome remodelling would be expected at these sites. To do this, we selected NFRs that are not bound by Abf1 or the GG-NER complex (n = 4415, a comparable number to the 3600 GCBS’s described earlier in Figure 3A). We observed no change in the nucleosome occupancy at these positions (Figure S3). By extension, relative rates of GG-NER at these sites would not be affected following deletion of *RAD16,* since these locations do not undergo GG-NER complex-mediated chromatin remodelling. Indeed, we observed that across a 2kbp window, repair rates are evenly distributed in both wild type and *rad16* mutant cells at this class of NFRs (see Figure 9B and D). In this case, the reduced repair rates in GG-NER defective cells are evenly distributed across these genomic positions. These observations confirm that the organisation of repair at GCBS’s is not a common feature of NFRs, but specifically requires the occupancy of the GG-NER complex, and the remodelling of its adjacent nucleosomes at these positions to initiate the GG-NER process.

## Discussion

In cells, maintaining the integrity of the genome is essential for life. Since DNA is constantly exposed to the deleterious effects of both the internal and external cellular environment, mechanisms have evolved to sense and repair the ensuing genetic damage. The ability to efficiently detect the presence of DNA damage that is packaged into chromatin is of paramount importance, and defects in the process are associated with a variety of diseases including cancer.

The main findings of this report reveal that chromatin remodelling during repair of DNA damage by nucleotide excision repair pathway (NER) is initiated from specific sites of GG-NER complex binding at the boundary sites of CIDs, which are genomic regions of higher-order nucleosome-nucleosome interaction. We show that in undamaged cells, the complex occupies these sites and is bounded by nucleosomes containing the histone variant H2A.Z. In response to DNA damage, we show that these boundary nucleosomes are evicted or exchanged in a GG-NER complex dependent fashion, and this enables the Rad7 and Rad16 components of the complex to redistribute to more distal sites within the CID. Finally, we demonstrate the importance of this mechanism to the removal of DNA damage by NER.

NER recognises and repairs a broad range of lesions, including those induced by UV light and a variety of chemical carcinogens. Two sub-pathways of NER exist that differ in their mechanism of initiating damage recognition. During transcription-coupled repair (TC-NER), recognition is initiated by the stalling of RNA polymerase II as it encounters the damaged DNA. This couples repair of DNA damage to the process of transcription and consequently to how this process is organised within the genome. This coupling results in an efficient mechanism for removing genetic damage and restoring gene expression to damaged transcribed DNA strands. Stalling of RNA pol II subsequently recruits NER factors that function in later stages of the NER process. These factors are common to later stage repair events in the global genome repair pathway (GG-NER). However, less is known about how repair of DNA damage is initiated and organised in the GG-NER sub-pathway, which repairs all non-transcribed regions of the genome. In yeast, this pathway relies upon a protein complex that is unique for the function of GG-NER. Using genomic techniques, we recently showed that GG-NER is organised into domains related to the promoter regions of open reading frames. We demonstrated that efficient DNA repair around these sites depends on the GG-NER complex regulating the histone acetylation status of nucleosomes in the vicinity, which alters the chromatin structure. However, until now, we have not established how chromatin remodeling is initiated during GG-NER by examining these events at the level of the nucleosome, the primary structural unit of chromatin.

To tackle this problem, we generated genome-wide nucleosome maps to analyse UV-induced changes to the nucleosome landscape. We analysed the genomic distribution of changes to the three core nucleosome parameters that quantify occupancy, fuzziness and position, to identify a subset of nucleosomes that are altered in response to UV-irradiation. These findings demonstrate that chromatin remodeling at this level occurs predominantly through dispersed local changes to nucleosome occupancy and fuzziness. However, nucleosome sliding in the context of gene expression can also be detected at known DNA-damage responsive genes, in line with previously published data (Lai and Pugh 2017). Our data shows that the remodelling of positioned nucleosomes adjacent to GCBS’s at many hundreds of genomic features in aggregate does not occur via nucleosome sliding. These findings are consistent with our previous biochemical observations (Yu et al. 2004; Yu et al. 2009). The complex can translocate across DNA *in vitro* through the activity of the SWI/SNF and helicase domains of Rad16, but it cannot slide nucleosomes *in vitro* (Yu et al. 2009). This mechanism may also drive the nucleosome remodelling events we describe here. Our current efforts are aimed at unraveling how the DNA translocase activity of the GG-NER complex contributes to the nucleosome remodelling demonstrated here. We show that the GG-NER complex binding sites identified in this study, are not simply regions of repair initiation, but are also locations of UV-induced histone eviction or exchange, involving nucleosomes containing the histone variant H2A.Z. We suggest that in undamaged cells these H2A.Z-containing nucleosomes represent a barrier that constrains and sequesters the GG-NER complex at these genomic regions. DNA repair may be initiated by removal of these barriers, allowing the GG-NER complex to redistribute from its initial binding locations. The loss of H2A.Z-containing nucleosomes removes the barrier for GG-NER complex redistribution. We suggest that this process might concurrently restrict RNA pol II transcription that requires H2A.Z-containing nucleosomes for efficient gene transcription (Weber et al. 2014). Therefore, this mechanism may contribute to the inhibition of bulk transcription in response to DNA damage, while at the same time driving the search for DNA damage by the GG-NER complex. Shut-down and restoration of normal gene expression is an established hallmark in maintaining the stability of the genome in response to DNA damage.

The occupancy of the GG-NER complex upstream of genes in undamaged cells, shows that it is an inherent component of chromatin, as well as playing a role in repairing its structure in response to damage. We speculate that the E3 ligase function of the complex is important in this process. Our previous work shows that the E3 ligase activity of the GG-NER complex is essential for binding of the complex to the chromatin (Yu et al. 2016). By extension, we expect that this activity is pivotal in maintaining the chromatin structure around GCBS’s in the absence of DNA damage. Future work will address this issue in order to understand how the E3-ligase activity maintains chromatin occupancy of the GG-NER complex, and to determine which factors or components of the chromatin are its targets.

Higher order chromatin structure in yeast has been identified following the introduction of methods that map distal nucleosome-nucleosome interactions, forming structural units that are classified as CIDs (Hsieh et al. 2015; Hsieh et al. 2016). These structures encompass 1 to 5 genes and range in size from a few kilobases up to 10Kbp. We find that ~50% of our GCBS’s can be found precisely at the boundaries between these genomic features. We speculate that the nucleosome-nucleosome interactions contained within CIDs may represent higher-order levels of structure that are remodelled during GG-NER. We plan to investigate this aspect in more detail in the future. We propose that the DNA translocase activity of the GG-NER complex could induce the remodelling of higher order chromatin structure, similar to the loop-extrusion model suggested for CTCF-Cohesin complexes in higher eukaryotes (Sanborn et al. 2015). In this model, two CTCF-Cohesin complexes bind to the chromatin and extrude DNA through the cohesin ring structure until they encounter a CTCF binding site (Sanborn et al. 2015). The CTCF and cohesin factors reside at the base or boundary of these loop structures, which may be analogous to the boundary positions to which the GCBS complex binds in the yeast genome. Although the loop-extrusion model has not been demonstrated in yeast, and the lack of a yeast homolog for CTCF, excludes the possibility of a direct parallel mechanism. However, we suggest that the redistribution of the GG-NER complex, by virtue of the DNA translocase activity of Rad16 could act as a wedge to disrupt the higher-order contacts that exist in the DNA loops that make up the CIDs. Future research aims to investigate the remodelling mechanism of higher-order chromatin structure using the micro-C methodology.

In summary, we conclude that our study demonstrates that in undamaged cells DNA repair complexes are positioned at hundreds of boundary regions that define the presence of CIDs; genomic domains of higher order nucleosome-nucleosome interactions. We suggest that this arrangement might represent origins of DNA repair initiation that promote the efficient repair of DNA damage in chromatin. We note that initiating chromatin remodelling from defined origins could effectively reduce the search space for DNA damage recognition, by compartmentalising the genome into functional modular chromatin structures that can be rapidly remodelled and efficiently repaired. Therefore, characteristic structural features of CIDs emerge when the genome is organised in this way – this ensures the rapid search and repair of genetic damage in chromatin.

## Methods

### Yeast strains used in this study

To delete the *RAD16* gene from the HA-Htz1 tagged W303-1B strain, we used a *RAD16::HIS3* disruption construct residing on pUC18. The pUC18 *RAD16::HIS3* (Reed et al. 1998) was digested using EcoRI and BamHI and used for transforming yeast. We used lithium-acetate transformation and selected for successful genomic integration of the disruption construct on His^-^ selection plates. From the transformation plate 12 individual colonies were restreaked on fresh media for single colony PCR and confirmation of UV phenotype. Successful clones were stored as glycerol stocks and used for the detection of genome-wide H2A.Z occupancy using ChIP-seq in the absence of *RAD16*.

### UV irradiation, yeast cell culture and crosslinking

Yeast cells were grown and UV irradiated as described previously (Yu, 2011, Yu 2016). Briefly, cells were grown to log-phase and resuspended in ice-cold 1x PBS to 2×10^7^ cells/ml. Using 10 J/m^2^s^1^, cells were irradiated with 100 J/m^2^ UV-C (254 nm). The cell suspension was kept in the dark to prevent photoreactivation and cells were spun down and resuspended in fresh YPD and incubated at 30°C to allow repair to take place. After the indicated repair time in YPD, cells were treated with formaldehyde to a final concentration of 1% to crosslink protein-DNA complexes for both ChIP-seq and MNase-seq analysis. The reaction was quenched in a final concentration of 125 mM Glycine. The cells were harvested by centrifugation and washed with ice-cold 1x PBS for 2 times. The final wash was performed using cold FA/SDS (50mM HEPES KOH pH 7.5, 150mM NaCl, 1mM EDTA, 1% Triton X-100, 0.1%, NaDeoxycholate, 0.1% SDS, 1mM PMSF) and the cells were transferred to a 2 mL Eppendorf tube. After pelleting the cells, they were snap-frozen in liquid nitrogen for long term storage.

### Chromatin preparation

Chromatin extracts were prepared as described previously (Teng, 2010, Yu 2011, Yu 2016). Briefly, cells were collected by centrifugation and prepared for lysis by bead-beating in FA/SDS (+PMSF) using 0.5 mL of glass beads for 10 minutes at 4°C. The whole cell extract was then sonicated with a Bioruptor (Diagenode) as described previously (Teng, 2010), after which the chromatin extract was collected by centrifugation. The chromatin extract is now ready for quantification and chromatin immunoprecipitation.

### Chromatin Immunoprecipitation

ChIP was performed as described previously (Powel 2015, Yu, 2011, Yu 2016). In summary, pre-washed Protein G Dynabeads™ (Invitrogen) were incubated with the optimal amount of monoclonal antibody (2 μg anti HA antibody Millipore, 05–904 or 3 μg anti Abf1 antibody yC20) in 50 μL of PBS-BSA (0.1%) for 30 min at 30°C in an Eppendorf thermomixer. The beads were then collected, washed and resuspended in 50 μL of PBS-BSA (0.1%) per sample and combined with 30 μL of 10x PBS-BSA (10mg/ml) and 300 μg of sonicated chromatin (~100-200 μL). The final volume was made up to 300 μL using PBS. The IP was performed at 21°C for 3 hours at 1300rpm in an Eppendorf thermomixer. Following this incubation, the samples were washed after which DNA was eluted in 125 μL of pronase buffer (25mM Tris pH 7.5, 5mM EDTA, 0.5% SDS) at 65°C at 900rpm for 30 min. Finally, pronase (Roche) was added to each sample and the cross-linking was reversed by incubating the samples at 65°C in a water bath overnight. Input samples containing 1/10th of IP sample were treated with pronase in parallel and incubated overnight as the IP samples. IP and input samples were treated with DNase-free RNase A (10mg/ml). For IP samples DNA was purified using the PureLink^TM^ Quick PCR Purification Kit (Invitrogen) and eluted with 50 μL elution buffer.

### Micrococcal nuclease digestion

MNase digest was performed as described previously (Kent). Briefly, yeast cells were UV treated as described earlier, except the final wash was not performed using FA/SDS but PBS. The pellet was resuspended in 250 μL 1 M Sorbitol for treatment with yeast lytic enzyme (YLE). YLE is prepared in a 200 μL volume containing 22.5 mg/mL YLE, 11.25 M 2-mercaptoethanol and 1 M sorbitol. The cells are treated for 3 min at room temperature. The cells were pelleted by pulse spin at 12,000 rcf in a microcentrifuge and washed carefully with 1 M sorbitol. The cells were pelleted again by pulse spin at 12,000 rcf, with the tubes now rotated 180°. After carefully removing the supernatant the cell pellet is ready for MNase treatment. Add 400 μL of digestion buffer containing 1 M sorbitol, 50 mM NaCl, 10 mM Tris-HCl (pH7.5), 5 mM MgCl2, 1 mM CaCl2, 1 mM 2-mercaptoehanol, 0.5 mM spermidine, 0.075% Nonidet P40, transfer to a 1.5 mL Eppendorf tube and add 4 μL of 30 U/μL MNase. Treat with MNase for 10 min at 37°C. The reaction can be terminated by adding a solution of 5% SDS and 250 μM EDTA. After RNase and pronase treatment the samples were incubated at 65°C overnight to reverse the protein-DNA cross-links. Finally, the DNA was recovered use phenol/chloroform extraction, ethanol precipitation and Invitrogen PCR kit purification, eluting the DNA in a final volume of 50 μL.

### Ion Proton Library preparation

This protocol was adapted from the LifeTechnologies *Ion ChIP-Seq Library Preparation on the Ion Proton^TM^ System,* Publication Number 4473623, Revision B. With minor modification to the blunt-ending, DNA purification and amplification as described below. We use ≤10 ng of ChIP DNA, quantified by using the Qubit® 2.0 Fluorometer. When samples are below the detection limit of the Qubit®, we used all of the ChIP DNA (40-50 µL).

### PreCR repair

Take 40 μL or ≤10ng of IP samples and 10 ng of Input samples diluted into 40 μL with ddH2O in a 1.5-mL Eppendorf LoBind® Tube and add the PreCR repair mixture (NEB). Incubate the reaction for 20 min at 37°C. Purify the DNA with magnetic beads using 1.8x sample volume of beads (CleanNA) according to the LifeTechnologies ChIP-seq protocol. It is important to use freshly (same day) prepared 70% ethanol for washing the beads for each purification. Elute in 50 μL low TE.

### End-repair & Blunt Ending

Use T4 DNA polymerase for end-repair and blunt ending. Combine 0.2 μL of T4 DNA polymerase (3,000 U/μL) in 1x NEB buffer 2 with 1.0 μL dNTPs (10mM) and 0.6 μL BSA (20 mg/ml) and 50 μL ChIP DNA in an end-volume of 120 μL. Mix by pipetting and incubate for 30 min at room temperature. Purify the DNA using magnetic beads using 1.8x sample volume of beads (CleanNA) according to the LifeTechnologies ChIP-seq protocol. Elute in 40 μL low TE. The DNA can be stored at 4°C at this point.

### Ligation

Prepare ligation mixtures using T4 DNA ligase (NEB, 400 U/μL) and the universal P1 adapter and A barcoded adapter and 10x Ligase buffer in an end-volume of 100 μL according to the LifeTechnologies ChIP-seq protocol. It is important to be careful and not to contaminate the barcoded adapters. Ligate for 30 min at room temperature. Note, this step is critical, longer incubations introduce adapter-adapter concatamers that get amplified during the next step. Purify the ligated DNA using magnetic beads using 1.5x sample volume of beads (CleanNA) according to the LifeTechnologies ChIP-seq protocol and elute in 40 µL low TE (the LifeTech protocol elutes in 20 µL). DNA can be stored at 4°C at this point.

### Nick Repair & Amplification

Use Q5 High Fidelity polymerase (NEB) to perform the nick repair reaction and amplification steps. These steps are performed in an end-volume of 100 µL. Combine 40 μL ChIP DNA with 2 units of Q5 HF polymerase and 1 μL forward and 1 μL reverse amplification primers (20 μM), 20 μL 5x Q5 reaction buffer, 2.5 μL dNTPs (10 mM), make up to 100 μL using ddH2O and mix by pipetting up and down. Place the tubes into a thermal cycler and run the following PCR cycling program (Table 1).

**Table 1.** Yeast strains and their respective genotype used in this study

| Yeast strain | Genotype |
| --- | --- |
| *BY4742 (WT)* | *MATα his3Δ1 leu2Δ0 lys2Δ0 ura3Δ0* |
| *Rad16-18myc* | *W303-1B Mat a RAD ade2-1 trp1-1 can1-100 leu2-3,112 his3-11,15 ura3-1* |
| *rad16Δ* | *W303-1B rad16Δ::HIS3* |
| *HA-HTZ1* | *W303-1B HA-HTZ1::KanMX* |
| *HA-HTZ1 rad16Δ* | *W303-1B HA-HTZ1::KanMX RAD16::HIS3* |

PCR Program:

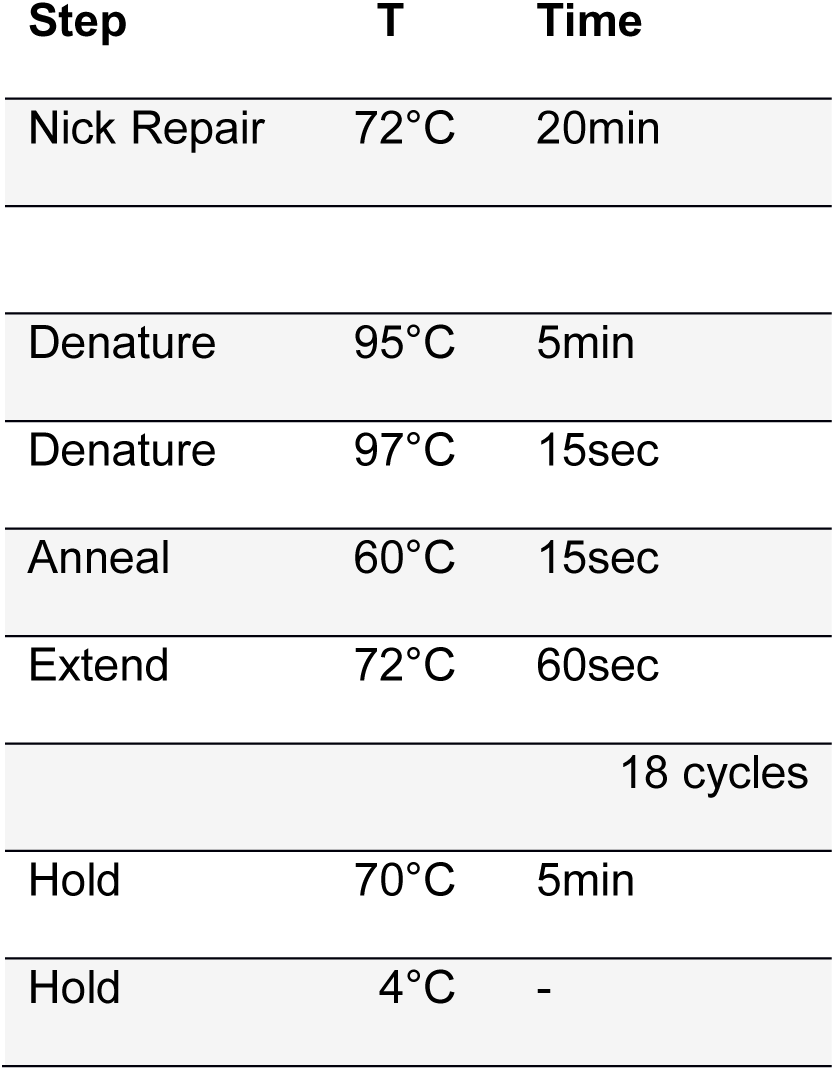

It is possible to optimise the number of PCR cycles if the input DNA is much lower than 1 ng. Purify with magnetic beads using 1.5x sample volume of beads (CleanNA) and keep a 2 µL aliquot before size selection for analysis on an TapeStation 2200 (Agilent Technologies) later.

### Size-selection and DNA purification using SPRI beads

Magnetic SPRI beads (CleanNA) are used according to the LifeTechnologies ChIP-seq protocol to size-select the library prep. The first round of purification uses a 0.7x sample volume of SPRI beads to selectively capture DNA >350bp, keeping the library DNA in the supernatant. During the second step 80 µL of SPRI beads are used (approx. 0.5x of sample volume), binding all DNA >160bp to the beads. Finally, the library DNA is eluted in 25 μL low TE.

### Quantification & Quality Control

Quality check the library prep before and after size selection by running the samples on a High Sensitivity tape on the TapeStation 2200 (Agilent Technologies). Quantify DNA using a 2 µL sample on the Qubit and convert to nM using:

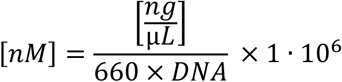

Prepare a dilution of 100pM of pooled libraries ready for sequencing. The emulsion PCR on the Ion Chef requires 50-100pM. At this stages sample were submitted for sequencing to our colleagues at the Wales Gene Park. The pooled library and individual library preps can be frozen at this point for long term storage.

### Next-generation sequencing using the Ion Proton platform

The pooled library is subjected to the Ion PI^TM^ Hi- QTM Chef Kit to generate Ion Sphere Particles (IPS) that are loaded onto the Ion PI^TM^ Chip v3 for sequencing on the Ion Proton Semiconductor Sequencer (ThermoFisher Scientific) according to the *Ion PI^TM^ Hi- Q^TM^ Chef Kit user guide*, publication number MAN0010967, revision B.0.

### Ion Proton oligonucleotide adapter sequences and annealing

The sequence and specifications of the universal P1 and barcoded A adapters were obtained from: *Ion Xpress^TM^ Plus gDNA and Amplicon Library Preparation* Publication Part Number 4471989 Rev. C, Revision Date 3 January 2012 appendix D. The oligos were ordered as ssDNA including the phosphorothioate bonds to protect from degradation and annealled manually. The complementary oligonucleotides were resuspended at equimolar 20 μM concentration in 1x TE complemented with 50 mM NaCl in an end-volume <500 μL. The mixture was heated to 95°C for 5 minutes in a heat block, after which the heat block was removed from the unit and allowed to cool to 25-30°C at room temperature (~1h30m). Correct annealing of a single product was confirmed by running a melt-curve protocol on a BioRad iCycler using 1 μL of template. All annealed oligos displayed a single peak in the melting curve around 78-80°C. The concentration was measured using a NanoDrop 2000 (ThermoFisher) and the adapters were stored at −20°C.

### Data Analysis

The trimmed Fastq files were aligned to the sacCer3 reference genome using the BWA-MEM 0.7.12-r1039 (Li and Durbin 2010) and piped into samtools 0.1.17 (r973:277) (Li et al. 2009) to convert the output to a sorted BAM file. These BAM files were used as input for downstream processing using MACS2 and DANPOS (see next section).

### Peak detection of ChIP-seq data using MACS2

In order to perform peak detection with MACS2 of two biological replicates, we merged the sorted bam files (input and IP) using samtools as suggested by the MACS2 developer notes before calling peaks. Running MACS2, we set the genome size to 12×10^6^, used a bandwidth of 100bp, allowed for a peak fold-change between 1 and 100 and set the regions that are checked around the peak positions to calculate the maximum local lambda to 2000 and 100000.

~~~
macs2 callpeak -t IP_merge.bam -c IN_merge.bam -f BAM -g 1.2e7 -n
merge -B --bw 100 -q 0.05 -m 1 100 --slocal 2000 --llocal 100000
~~~

MACS2 outputs the normalised input and IP traces as bedgraph files and the peak and summit position as tab-delimited data that were loaded in IGB (Freese et al. 2016) for inspection. The peaks called for Abf1 binding in this output were used for the annotation and overlap calculations used to characterise the GCBS’s described in the manuscript.

### Nucleosomes Mapping of MNase-seq data using DANPOS

Aligned MNase-seq data, as sorted bam file format, was submitted to DANPOS for mapping nucleosome positions. Submitting multiple datasets to DANPOS allows for fold-change normalisation that compensates for different read-depth or coverage between datasets. Using the MNase-seq data from wild type and *rad16* deleted cells from non-irradiated (-UV) and 0 minutes and 30 minutes post-UV samples, DANPOS successfully and consistently maps ~65,000 nucleosomes for each dataset. DANPOS outputs the statistical information on position, fuzziness and occupancy in a spread-sheet format and produces a wig-file that contains the genome-wide trace of the nucleosomes positions. These wig files were accessed through IGB (Freese et al. 2016) for viewing and generating snapshots included in the manuscript. This output was also submitted to SeqPlots (Stempor and Ahringer 2016) for plotting (see following section). All composite plots described in the manuscript used the data described here.

### GCBS annotation using ChIPpeakAnno

The Abf1 binding sites as detected by MACS2 were loaded in the R-statistical environment and annotated using the ChIPpeakAnno R-package (Zhu et al. 2010). BED files containing the coordinates of the Abf1 binding sites, genome-wide NFRs (Yadon et al. 2010) and Abf1 consensus motifs (Bailey and Elkan 1994; Khan et al. 2018) were loaded using the *BED2RangedData* function. *makeVennDiagram* was used to generate the Venn diagram as shown in Figure 3. The final Venn diagram as displayed in the figure was scaled in photoshop to roughly represent the number of genomic intervals of each feature.

Using *annotatePeakInBatch*, ensembl annotation data was used to map the GCBS’s to the nearest genomic features such as TSS or Gene. *AnnotatePeakInBatch* provides every genomic interval that represents a GCBS with an orientation to the nearest gene as upstream, overlapStart, downstream, overlapEnd, inside or includeFeature. The Abf1 binding sites that represent the GCBS’s are predominantly of the upstream and overlapStart class. We used this annotation data to specifically select this class of Abf1 binding sites and used the information of these associated genes to retrieve their TSS coordinates to make the ORF plots presented in the manuscript.

### Visualisation of genome-wide ChIP-seq, MNase-seq and ChIP-chip data

Genomic intervals of binding sites, NFRs, motifs or ORFs can be submitted to SeqPlots as canonical BED files. Genome-wide traces of continuous data such as those of MNase-seq and ChIP-seq data can be uploaded as wig or bigWig (.bw) files. To convert ChIP-chip and 3D-DIP-chip data into a format that was amenable for plotting we converted the output from Sandcastle (Bennett et al. 2015) the following way. Using the *writeCCT* script from Sandcastle we export the array data in a tab-delimited format. For compatibility we converted the chromosome names to roman numerals using standard command line operations using Perl. This rudimentary BED file can now be converted to a wig file using the UCSC *BedToWig* script (http://genomewiki.ucsc.edu/index.php/File:BedToWig.sh) setting the span to 100. Finally, the wig-file was manually converted to a bigWig file using the UCSC wigToBigWig script (http://hgdownload.cse.ucsc.edu/admin/exe/linux.x86_64/wigToBigWig) using a file containing the chromosome sizes from sacCer3 in the process. Some features overlapped and had to be manually corrected. Data that was loaded into SeqPlots was navigated using the SeqPlots gui as per the authors instructions. All figures presented in this manuscript were generated using SeqPlots, exported and processed using Adobe Illustrator and Photoshop.

### Micro-C data processing and plotting

Datasets from the Micro-C XL experiments were retrieved from the ENA repository (Study PRJNA336566). From the double cross-linked data available we opted for the 3% FA and 3mM DSG for 40 minute dataset (SRR4000672) for plotting figure 5. We use HiC pro to align the data, build the contact maps, normalise the data and QC according to the authors documentation. HiC pro can handle data generated from MNase treated chromatin as opposed to many other packages that rely on restriction enzyme fragmented DNA to map HiC interaction. Next we used HiC-plotter to visualise the Micro-C XL data in conjunction with our MNase-seq and ChIP-seq data.

### Data access

We obtained the HiC boundary positions from the supplementary data accompanying the Hsieh et al. Manuscript (Hsieh et al. 2015). A list of genome-wide NFRs was obtained from Yadon et al. (Yadon et al. 2010). The list of Abf1 consensus motifs can be obtained using the MEME fimo algorithm. The data described in this manuscript was submitted to ArrayExpress (http://www.ebi.ac.uk/arrayexpress) and can be retrieve using accession code E-MTAB-6569.

## Acknowledgements

We would like to acknowledge our colleagues at the Wales Gene Park for their technical and bioinformatic support in generating the NGS data. We would like to thank Stephen P. Jackson for critically reading the manuscript. The work was supported by an MRC research grant MR/K000926/1 to SHR.

## Author Contributions

SHR and PVE conceived of the study and experiments. PVE and SPN performed the wetlab experiments. PVE analysed the data. SHR and PVE interpreted the data and wrote the manuscript. The other authors contributed ChIP-Chip or 3D-DIP-Chip experiments.

## Disclosure Declaration

The authors declare no conflict of interest

## Supplementary Figures

**Figure S1.**
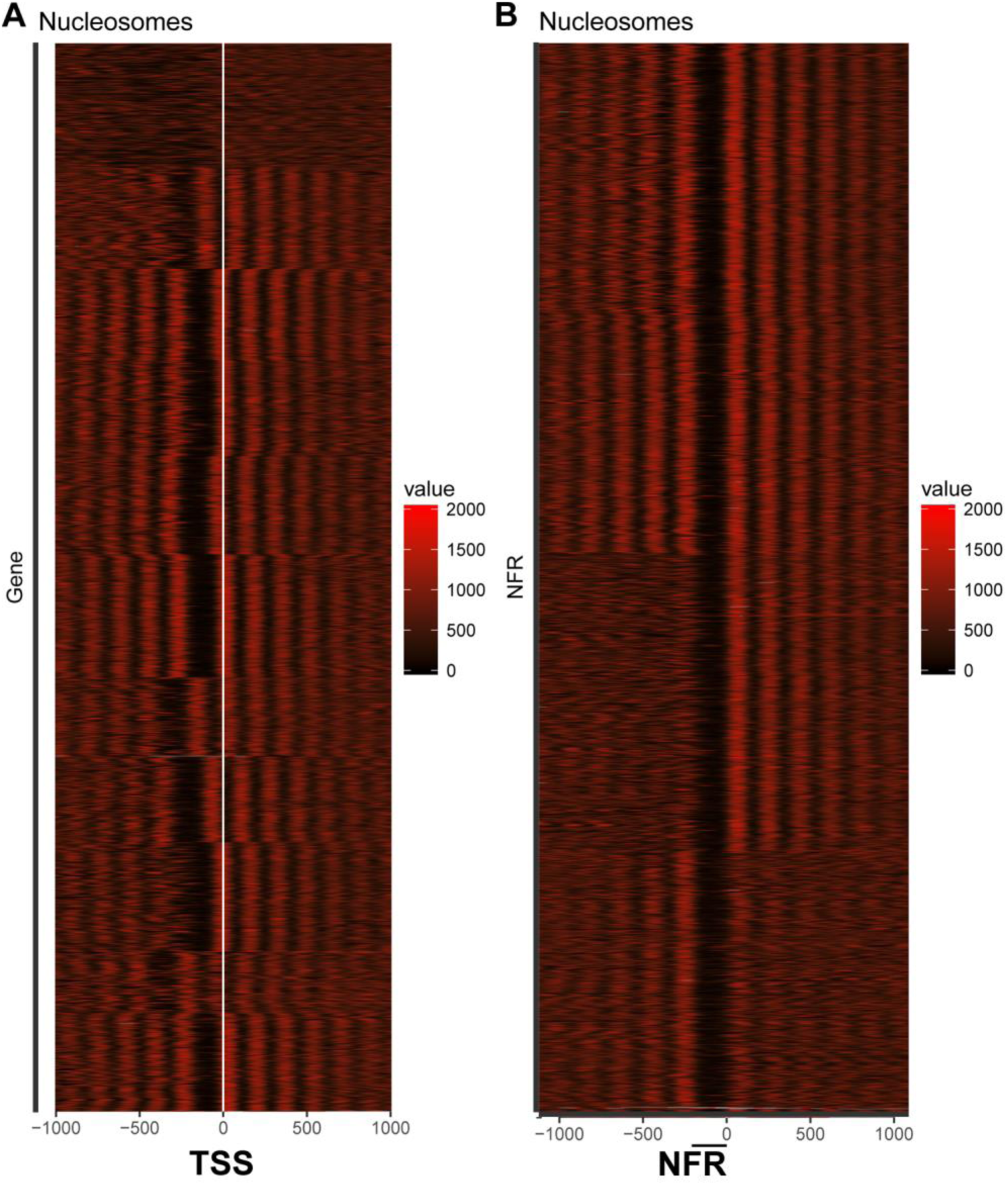
The relationship of TSS to NFR distance and +1 nucleosomes at genes downstream of GCBS’s are displayed here. Heatmaps were generated using Seqplots and transferred to the statistical R-environment for further analysis. A) Using K-means clustering analysis, the ~2600 genes that are annotated to the GCBS’s, were grouped into 13 clusters of similar structure. The heatmap displays the relative nucleosome density around the TSS highlighted with the with line in the middle, including 1 Kbp up- and downstream. The intensity of the heatmap is proportional to the normalised read-depth indicated in the legend. B) The same nucleosome data as displayed in A) was used but now aligned to the right side of the accompanying NFR. K-means clustering of the data in this orientation resulted in the delineation of 4 clusters. The legend indicates the intensity of the heatmap as a function of normalised reads. The bar in the bottom indicates the orientation of the NFR at these genomic positions.

**Figure S2.**
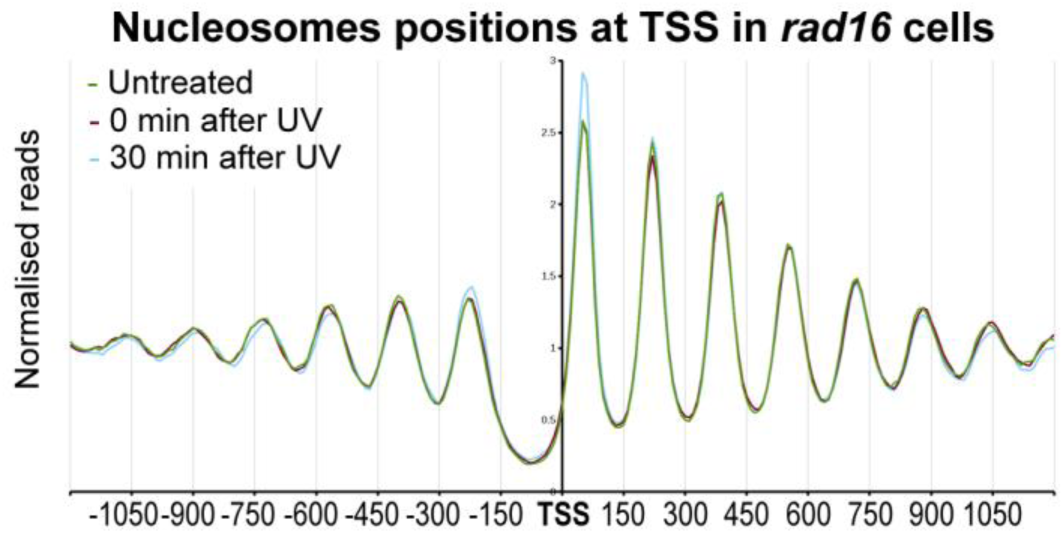
Nucleosomes occupancy around all TSS’s in yeast does not change in response to UV irradiation in *rad16* cells. Composite plot of nucleosomes positions relative to all TSS (n = 5,171). Genome-wide MNase-seq data was used to aggregate nucleosome positioning in relation to TSS positions.

**Figure S3.**
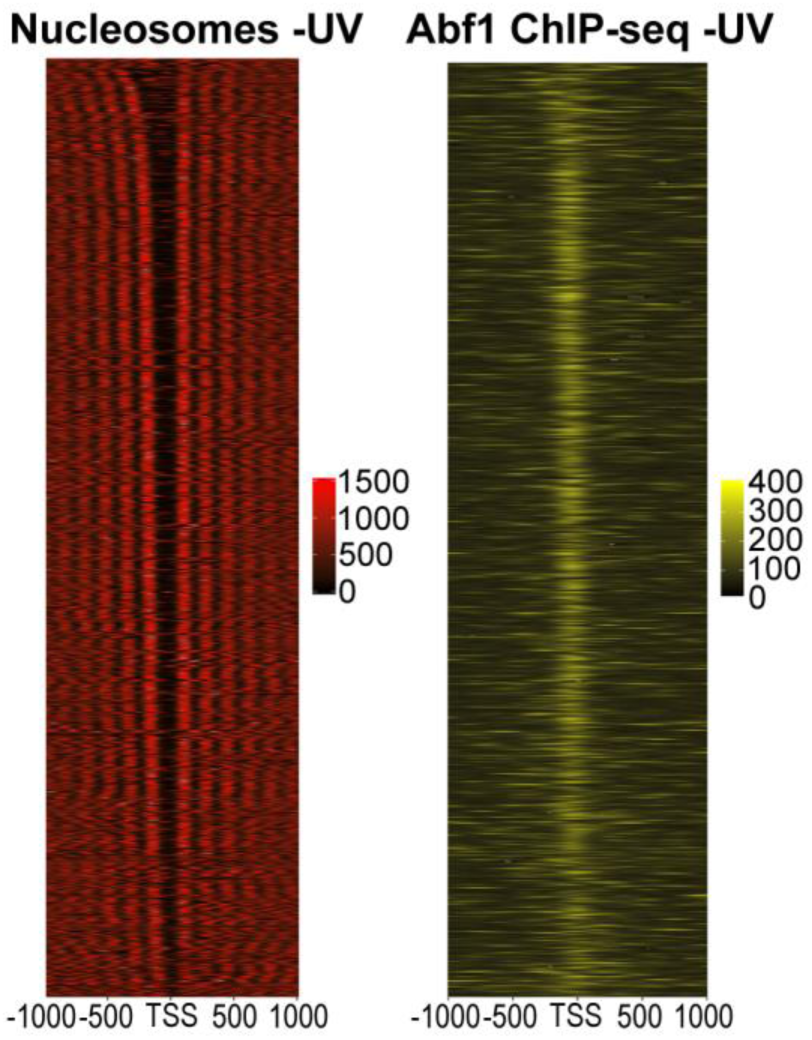
Nucleosome positions and Abf1 binding to chromatin in relation to TSS is linked to NFR width. Nucleosome data and Abf1 ChIP-seq data from Figure 8A are plotted at GCBS-associated TSS’s as a heatmap. The intensity is a measure of occupancy in reads indicated to the right of each heatmap. Plots were generated using SeqPlots and transferred into the R-statistical environment. Here the nucleosome traces were sorted by NFR-size from widest (top) to narrowest (bottom).

**Figure S4.**
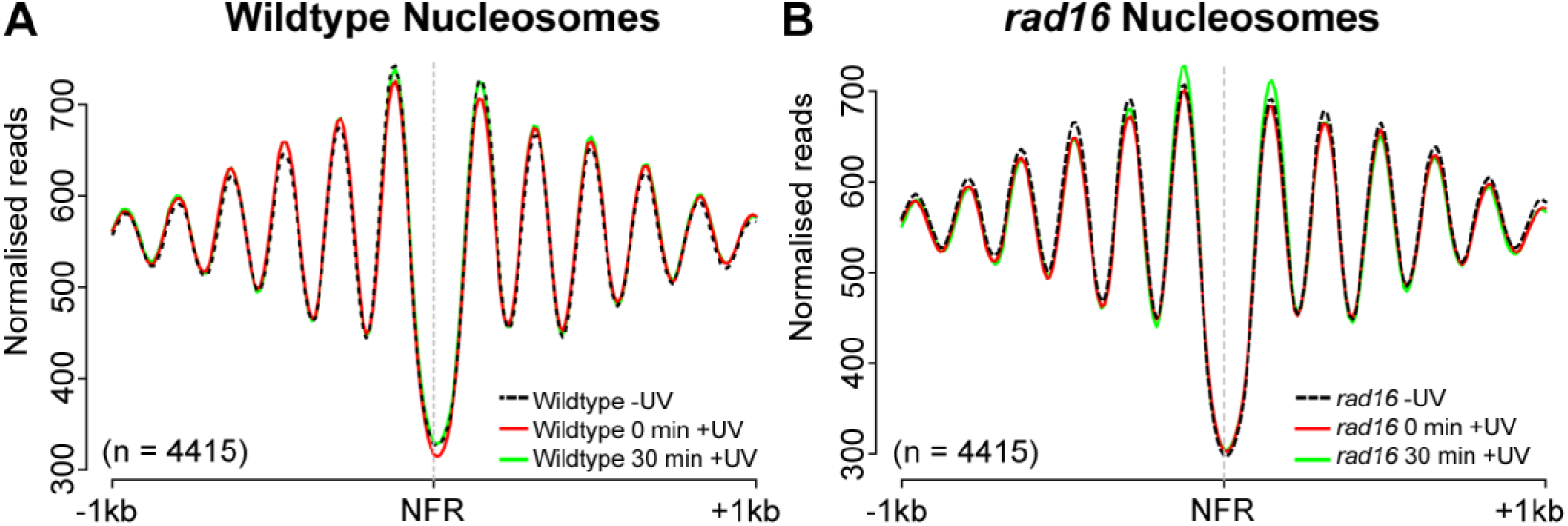
Nucleosome positioning around non-GCBS-bound NFRs in wild type and *rad16* mutant cells do not change in response to UV irradiation. The NFR positions (n = 4415) that do not overlap with an Abf1 binding sites or Abf1 consensus motif, as described in Figure 3A, were used to plot the MNase-seq nucleosome data as composite plots in wild type (A) and *rad16* deleted cells (B). On the x-axis the NFR position and 1 Kbp regions up- and downstream are displayed, while the y-axis depicts the normalised reads to indicate the nucleosome occupancy as these genomic locations.

